# CK2 variant function and disease modelling in *Drosophila* reveal allelic heterogeneity and Wnt/β-catenin-mediated phenotypes

**DOI:** 10.64898/2026.08.20.746075

**Authors:** Yina Her, Danielle M. Pascual, Ying Lao, Harsimran Kaur, Alondra Griffiths, Robert Beattie, Bradley W. Doble, Patrick Frosk, Rene P. Zahedi, Paul C. Marcogliese

## Abstract

Heterozygous pathogenic variants in *CSNK2A1* or *CSNK2B* encoding the Casein Kinase 2 (CK2) protein complex, lead to pediatric neurodevelopmental disorders, Okur-Chung Neurodevelopmental Syndrome (OCNDS) and Poirier-Bienvenu Neurodevelopmental Syndrome (POBINDS). OCNDS and POBINDS are characterized by a range of symptoms, including developmental delay, intellectual disability, facial dysmorphism, and seizures. Despite over 250 reported cases of OCNDS and POBINDS, we do not fully understand how specific alterations in CK2 relate to the heterogeneity observed in patients. To investigate this, we used the fruit fly, *Drosophila melanogaster*, as a model system. To assess variant impact, we co-expressed human *CSNK2A1* and *CSNK2B* reference or disease-causing variants in flies. In parallel, we determined the role of *Drosophila CkII* in the developing and mature nervous system, specifically in neurons and glia. We found that 12/13 variants tested act as full or partial loss-of-function with one *CSNK2A1* variant showing gain-of-function. Phospho-proteomic studies in neurons revealed separate signatures for loss- and gain-of-function variants. We found that neuronal and glial CkII is critical for organismal development. Reduction of neuronal *CkII* in the adult nervous system causes motor and seizure-like phenotypes. Finally, given the known role of CK2 in potentiating Wnt/β-catenin signalling, we show that Wnt agonists partially rescue phenotypes associated with adult-specific neuronal reduction of *CkII*. This work generates *Drosophila* models of *CSNK2A1* and *CSNK2B* expression to functionally assess variant impact, as well as an adult-specific neuronal loss-of-function model for drug screening and mechanistic studies.

## Introduction

Okur-Chung Neurodevelopmental Disorder (OCNDS) (1) and Poirier-Bienvenu Neurodevelopmental Disorder (POBINDS) (2) are rare pediatric neurodevelopmental disorders (NDDs). Clinically, both NDDs are characterized by global developmental delay, intellectual disability, motor issues, behavioural abnormalities, seizures, and dysmorphic facial features. Although over 250 affected individuals have been reported, there is a widening spectrum of phenotypes reflecting the heterogeneity of these NDDs (3,4). OCNDS and POBINDS are primarily caused by *de novo* pathogenic variants in *CSNK2A1* or *CSNK2B* which encode the catalytic (CK2α) and regulatory (CK2β) subunits of Casein Kinase 2 (CK2), respectively. CK2 is ubiquitously expressed and enriched in the nervous system (5). It is highly conserved across eukaryotes and is constitutively active as a serine/threonine kinase with more than 300 known targets (6–8). CK2 has been implicated in multiple cellular pathways as a key regulator for essential cellular functions such as growth, differentiation, and signalling (9). The proposed mechanism of disease for OCNDS and POBINDS is presumed haploinsufficiency of either subunit (1,10). *In vitro* experiments have shown that some pathogenic variants decrease CK2 stability, holoenzyme assembly, and kinase activity (11,12). Although one patient-derived *Csnk2a1* p.K198R knock-in mouse model has been characterized (13), no other *in vivo* models harbouring multiple CK2 pathogenic variants have been reported to date.

*Drosophila melanogaster*, commonly known as fruit flies, are a powerful *in vivo* model to investigate the functional assessment of patient-specific CK2 coding variants. Increasingly, flies have been used as a living system to assess the functional impact of human variants in NDDs (14,15). In this study, we use the well-established *GAL4/UAS* system to compare the functional impact of CK2 reference and NDD-related variant expression in a ubiquitous and tissue-specific manner. By examining morphological and phospho-proteomic impact, we found that most CK2 pathogenic variants act as full or partial loss-of-function (LoF or pLoF), with the CSNK2A1 p.R47Q variant acting as a gain-of-function (GoF). RNA interference (RNAi) -mediated pan-neuronal or pan-glial knockdown of *Drosophila CkIIα* or *CkIIβ* during development causes lethality. Hence, we determined the role of *CkIIα* and *CkIIβ* in adult neurons and glia using the GeneSwitch system (16). We show that reduction of *CkIIα* or *CkIIβ* in adult neurons causes motor and seizure-like behaviour that can be partially ameliorated by enhancing Wnt/β-catenin signalling. Behavioural phenotypes were not observed upon adult-specific glial knockdown. By integrating variant-specific modelling with nervous system knockdown studies, this work reveals functional heterogeneity in CK2-related NDDs and points to Wnt/β-catenin signalling as a potential key mediator of LoF phenotypes.

## Results

### Co-expression of *CSNK2A1* and *CSNK2B* causes overexpression phenotypes in flies

Given that the CK2 holoenzyme is constitutively active (6), we first determined the impact of ubiquitous expression of reference single genes *CSNK2A1* or *CSNK2B*, or tandem co-expression of both cDNAs: *UAS* (*Upstream Activation Sequence*) *- CSNK2A1::T2A::CSNK2B* - hereafter referred to as *CK2^REF^*. Ubiquitous overexpression of *CK2^REF^* using *actin-GAL4* results in no viable adults, causing lethality at 25°C, in both male (**Fig. 1A**) and female flies (**Fig. 1B**). This contrasts with the expression of single cDNA expression of individual CK2 subunits (*CSNK2A1*, *CSNK2B*, or *CSNK2A2* - the latter not connected to disease), resulting in viable flies. This suggests that co-expression of both human a and b subunits produces functional outcomes in flies. To gain deeper insight, we chose to express UAS-cDNA constructs in the developing wing pouch with *nub-GAL4*. The developing *Drosophila* wing provides a robust platform for expressing human cDNA, as it is largely dispensable for organismal viability and allows for precise morphological analysis within a well-characterized signalling landscape that parallels Wnt-dependent neural development (17,18). Wing pouch expression of *CK2^REF^*causes decreased wing size in male (**Fig. 1C**) and female (**Fig. 1D**) flies at 29°C. The wings of *nub-GAL4 > CK2^REF^* flies also show abnormal morphological phenotypes, including blisters as well as loss of anterior and posterior crossveins (**Fig. 1E**). Individual CK2 subunits reduce wing size more modestly than *CK2^REF^*, and they do not produce *CK2^REF^*-associated morphological abnormalities. These data suggest that co-expression of *CSNK2A1* and *CSNK2B* inhibits wing development and can be utilized as a readout to assess variant impact.

**Figure 1:**
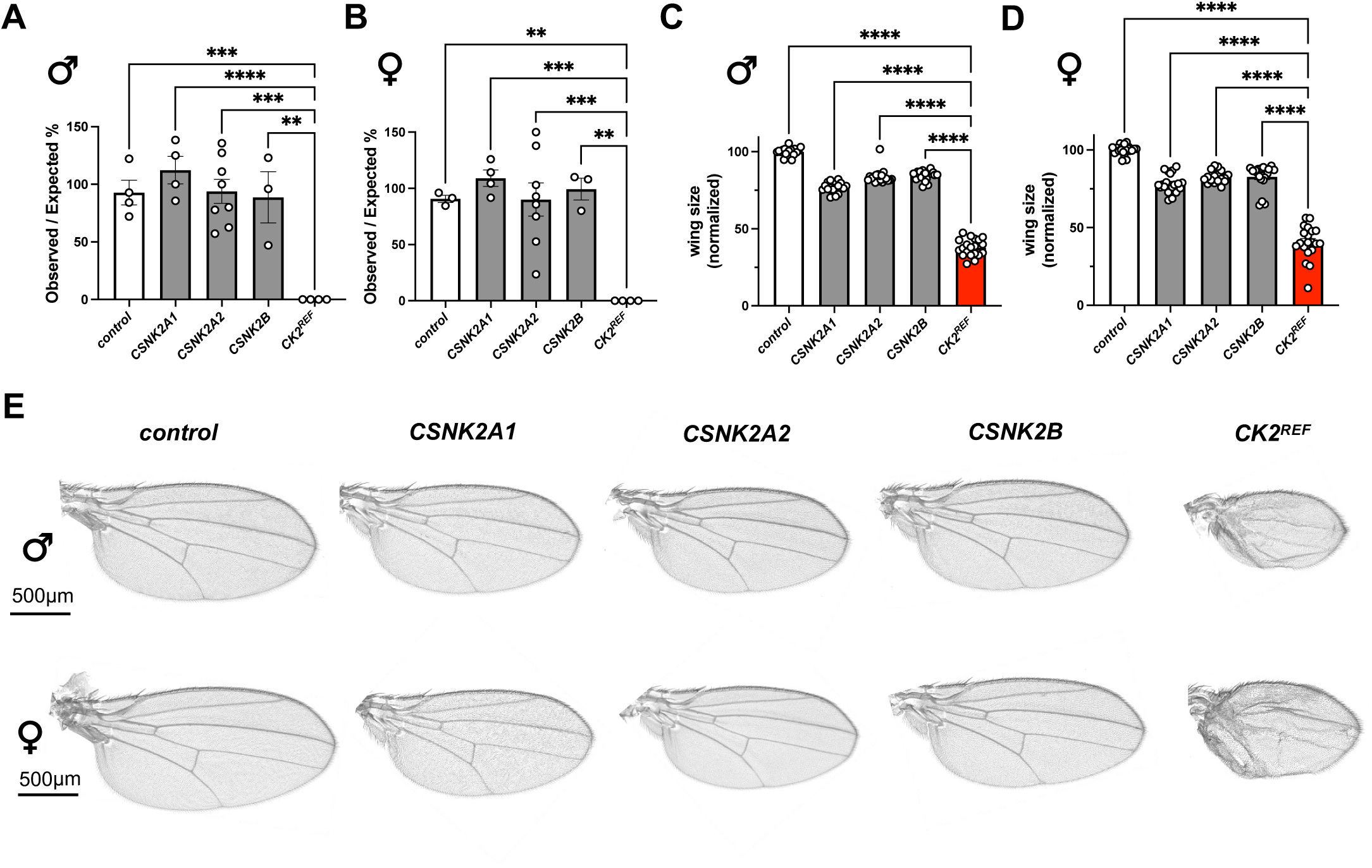
Tandem co-expression of *CSNK2A1* and *CSNK2B* causes overexpression phenotypes in flies. (A & B) Mendelian ratios of viable adult flies upon single gene or tandem co-expression of *CSNK2A1* and *CSNK2B* (*UAS-CSNK2A1::T2A::CSNK2B*; herein referred to as *CK2^REF^*) using *actin-GAL4* at 25°C in males (A) and females (B). (C & D) Single gene or *CK2^REF^* expression in the developing wing pouch using *nub-GAL4* at 29°C in males (C) and females (D). (E) Representative optical images of adult wings. ANOVA followed by Tukey’s post hoc test, **p<0.01, ***p<0.001, ****p<0.0001. Data were derived from three independent genetic crosses for statistical analysis.

### Multiple assays in *Drosophila* reveal a spectrum of full and partial loss-of-function and a gain-of-function for OCNDS and POBINDS variants

To determine the functional impact of OCNDS and POBINDS variants *in vivo*, we generated 13 pathogenic variants in the *CK2^REF^* construct and transgenic UAS-cDNA flies. Selected missense variants were prioritized based on hotspots, present in multiple reports (1,19–25), and interspersed locations throughout the proteins (**Fig 2A and B**). Critically, all UAS-cDNA constructs are inserted into the same genomic location in the same strain, controlling for positional effects and allowing for direct functional comparison of *CK2^REF^* to the variants. Protein expression was confirmed by western blot in flies expressing CK2 cDNA constructs pan-neuronally using *neuronal Synaptobrevin* (*nSyb*)*-GAL4* (**Fig. 2C-H**). Although ϕC31-mediated transgenesis allows for similar UAS-cDNA expression, the CSNK2A1 variants p.Y50C, p.K198R, and p.R312W showed decreased CSNK2A1 protein levels (**Fig. 2D**). CSNK2A1 p.K198R, and p.R312W also resulted in lower CSNK2B protein levels, albeit with an antibody that cross-reacts with *Drosophila* CkIIβ (**Fig. 2E**). This suggests altered stability. Similarly, CSNK2B variants p.D32N and p.C137R showed lower protein levels (**Fig. 2G and H**). The ubiquitous expression of *CK2^REF^* results in lethality when using *actin-GAL4* at 25°C (**Fig. 2I and J**). The ubiquitous expression of three missense *CSNK2A1* variants (p.R47Q, p.R80C, and p.K198R) results in lethality, similar to *CK2^REF^* (**Fig. 2I**). This suggests that these variants retain some function in this assay. Four *CSNK2A1* missense variants, p.Y50C, p.S51R, p.D175G, and R312W, display complete viability (**Fig. 2I**), suggesting strong LoF in this assay. For *CSNK2B*, the ubiquitous expression of one truncation variant p.W9* and the p.C137R missense variant results in viable flies (**Fig. 2J**), indicating strong LoF. Three missense variants (p.D32N, p.R86C, p.R111P) show complete lethality, indicative of similar function to *CK2^REF^* in this assay (**Fig. 2J**). Finally, the p.H165R shows lethality in males, indicating no LoF, whereas flies are partially viable in females, suggesting partial LoF (**Fig. 2J**). Together, this data suggests variable function across pathogenic variants.

**Figure 2:**
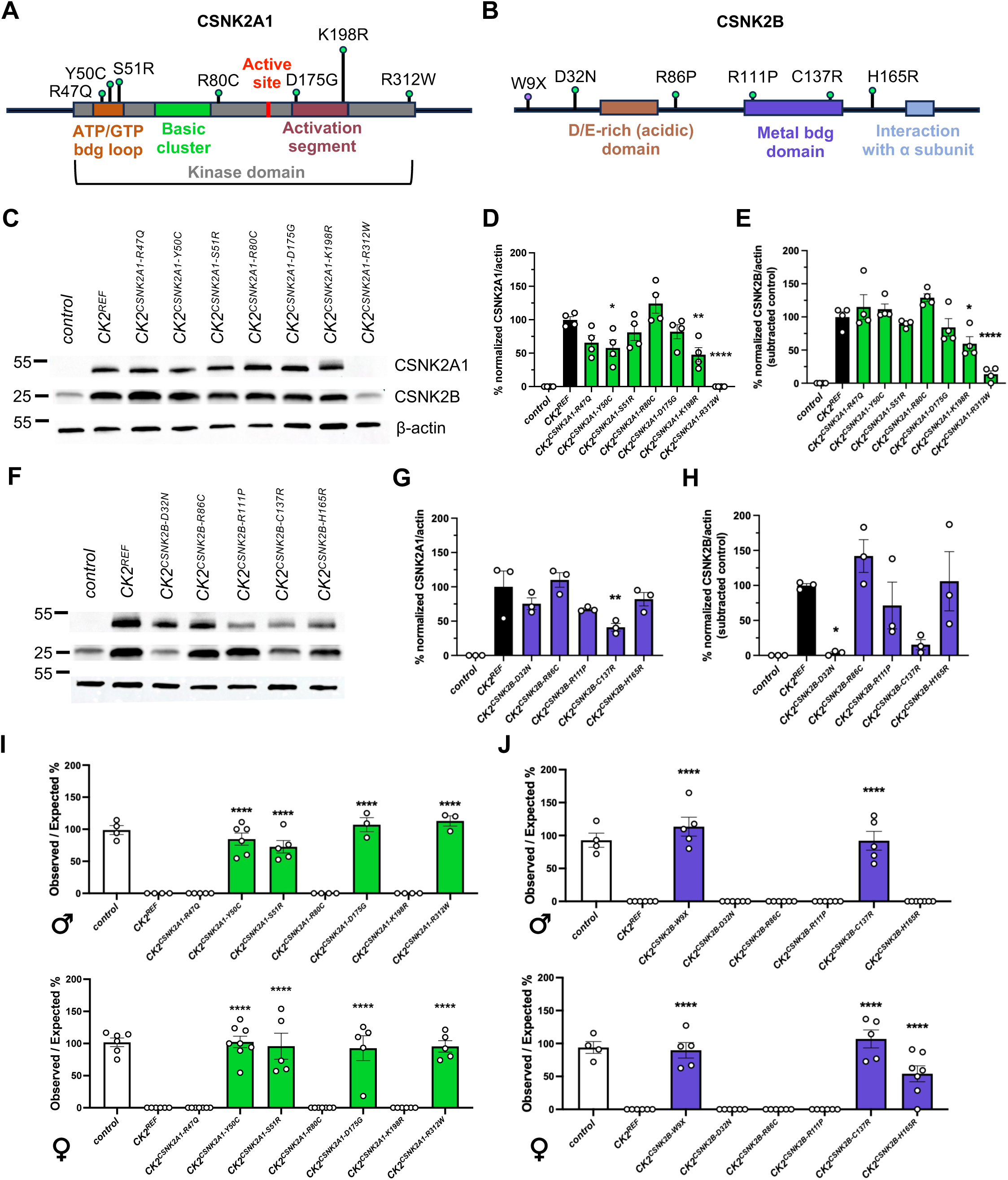
Protein structure, quantification, and function of pathogenic *CSNK2A1* or *CSNK2B* variants expressed in flies reveals allelic heterogeneity. (A and B) Schematic of CSNK2A1 and CSNK2B with conserved domains. (C-H) Protein expression and quantification of CK2 in neurons using *nSyb-GAL4* by western blot. (I and J) Ubiquitous overexpression of *UAS-CSNK2A1* and *UAS-CSNK2B* variants using *actin-GAL4* at 25°C in males and females. ANOVA followed by Tukey’s post hoc test, *p<0.05, **p<0.01, ***p<0.001, ****p<0.0001. Mendelian data was derived from a minimum of four independent genetic crosses with a minimum of 20 flies analyzed per sex and per cross.

The expression of *CK2^REF^* in the developing wing pouch using the *nub-GAL4* at 29°C causes decreased wing size and highly penetrant blisters (**Fig. 1E and Fig. 3A and E, Supplementary Material, Fig. S1A and E**). These morphological abnormalities are not present in control wings. For *CSNK2A1*, all seven missense variants showed a significant increase in wing size compared to *CK2^REF^* expression (**Fig. 3B, Supplementary Material, Fig. S1B**), suggesting some level of LoF for all pathogenic variants. Four missense variants, p.Y50C, p.S51R, p.D175G, and p.R312W, failed to produce a strong blister phenotype compared to *CK2^REF^* (**Fig. 3D, Supplementary Material, Fig. S1D**), suggesting strong LoF. The p.R47Q, p.R80C, and p.K198R showed no significant changes in the presence of blisters in comparison to *CK2^REF^* (**Fig. 3D**). This suggests that the *CSNK2A1* p.R47Q, p.R80C, and p.K198R variants retain some function and are likely pLoF. All *CSNK2A1* variants tested, except the p.R47Q, did not cause any bristle loss phenotypes along the wing margin when compared to *CK2^REF^* (**Supplementary Material, Fig. S2A-B**). Although the p.R47Q variant showed some pLoF regarding wing size, these wings also consistently showed the presence of ectopic hairs, that were absent from *CK2^REF^* or any other variant examined in this study (**Fig. 3C, Supplementary Material, Fig. S1C**). This suggests potential GoF activity for p.R47Q, specifically acting as a neomorphic allele.

**Figure 3:**
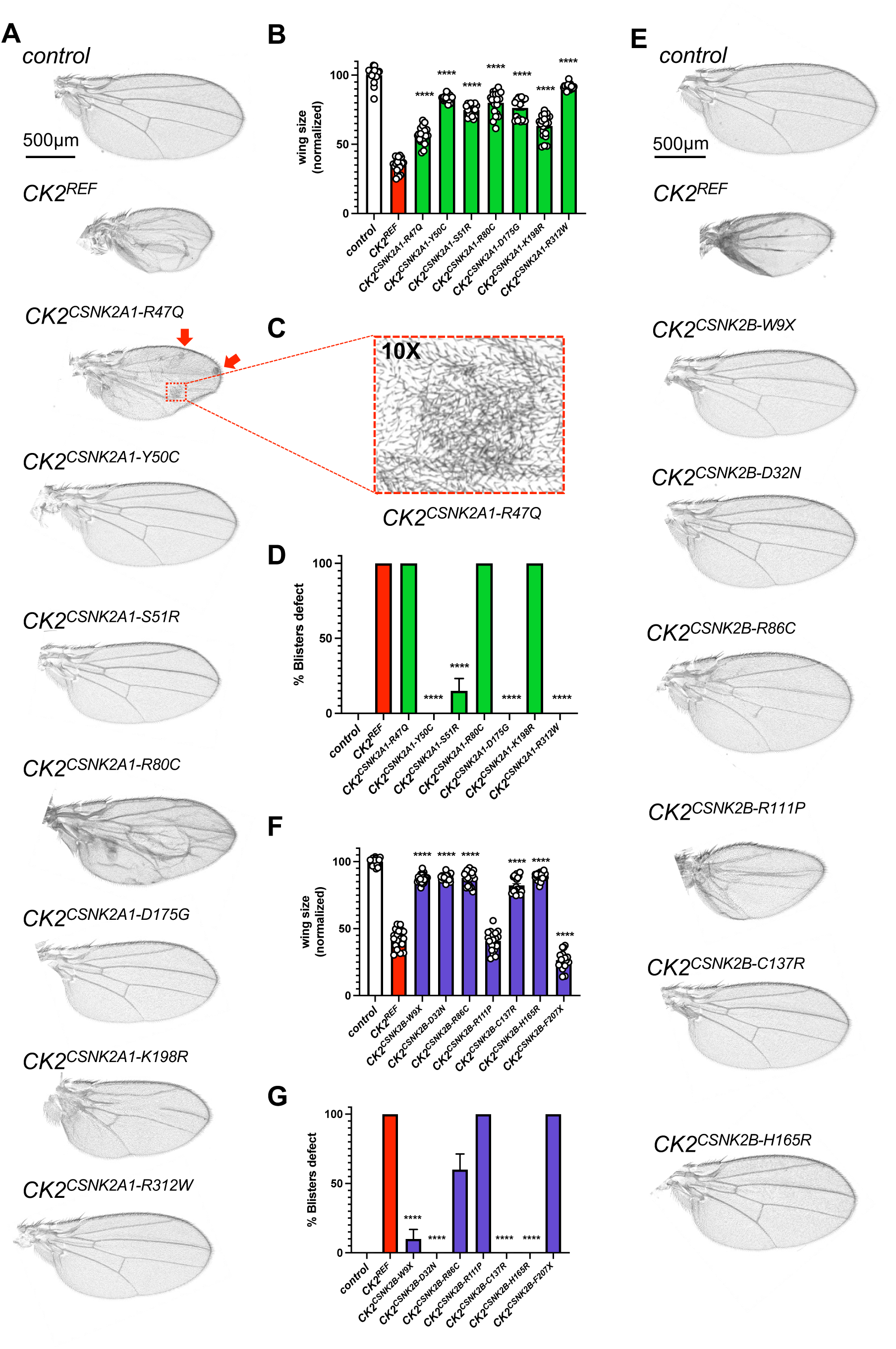
Wing pouch expression of pathogenic *CSNK2A1* or *CSNK2B* variants reveals a spectrum of LoF with the CSNK2A1 p.R47Q acting as a GoF. Representative wing images upon wing-specific expression of *UAS-CSNK2A1* variants (A) and *UAS-CSNK2B* (E) using *nub-GAL4* at 29°C. Quantification of wing size and presence of blisters for *UAS-CSNK2A1* (B & D), *UAS-CSNK2B* (F & G). (C) 10X zoom image of the CSNK2A1 p.R47Q variant showing ectopic hair phenotype. (B & F) ANOVA followed by Tukey’s post hoc test, ****p<0.0001. (D & G) Kruskal-Wallis test followed by Dunn’s multiple comparisons test, was completed with significance defined as *p<0.05, **p<0.01, ***p<0.001, ****p<0.0001. Data were derived from three independent genetic crosses, with a minimum of n=20 in each cross. All data derived from male flies (see supplement for female data).

For *CSNK2B*, five variants (p.W9*, p.D32N, p.R86C, p.C137R, and p.H165R) showed increased wing size compared to *CK2^REF^* (**Fig. 3F, Supplementary Material, Fig. S1F**). This suggests these variants act as LoF. The p.R111P did not show any significant difference in wing size to *CK2^REF^*, suggesting similar function. The p.W9*, p.D32N, p.C137R, and p.H165R do not produce significant blisters in comparison to *CK2^REF^* (**Fig. 3G, Supplementary Material, Fig. S1G**). Again, these data suggest LoF. Two missense variants, p.R86C and p.R111P, show a consistent blister phenotype like *CK2^REF^*, suggesting some partial function in this assay. Given that the p.R111P acted like *CK2^REF^* in wing size and blisters, we examined wings for any bristle loss along the wing margin. Expression of *CK2^REF^* with *nub-GAL4* produces bristle loss in 100% of wings examined, as the p.R111P as well as all other *CSNK2B* variants fail to cause bristle loss (**Supplementary Material, Fig. S2C-D**). Taken together, these results suggest that the p.W9*, p.D32N, p.C137R, and p.H165R act as strong LoF, as the p.R86C acts as a pLoF with the p.R111P variant acting as a relatively milder pLoF in these assays.

### Phospho-proteomic neuronal profiling shows differential targets for key variants *in vivo*

To determine whether the functional classifications obtained by tissue-specific overexpression assays reflect changes in CK2 kinase activity in the *Drosophila* nervous system, we performed quantitative phospho-proteomics. We selected three representative variants based on their distinct behaviours: two strong LoF variants, one for each gene (*CK2^CSNK2A1^ ^p.D175G^* and *CK2^CSNK2B^ ^p.W9X^*), and one variant that exhibited GoF/neomorphic activity in the wing (*CK2^CSNK2A1^ ^p.R47Q^*). Pan-neuronal expression of *UAS-CK2^REF^* using *nSyb-GAL4,* significantly increased phosphorylation of multiple known CK2 target sites compared to the *UAS-empty* control (**Fig. 4A**). By contrast, expression of either of the LoF variants (*CK2^CSNK2A1^ ^p.D175G^* or *CK2^CSNK2B^ ^p.W9X^*) showed markedly reduced phosphorylation at these same top sites relative to *CK2^REF^*(**Fig. 4B and C**). The heatmap highlights the top *CK2^REF^*-upregulated phospho-sites and confirms their reduction with both LoF variants (**Fig. 4E**). These data suggest that patient-derived LoF variants impair CK2 kinase activity. Although the neomorphic p.R47Q variant also showed a reduction in phosphorylation of these top targets compared to *CK2^REF^* (**Fig. 4D and E**), the p.R47Q variant also showed specific phospho-targets that were absent from *CK2^REF^*or the LoF variants (**Fig. 4F**). These novel phospho-targets included proteins: Nak, Fife, Raf, RhoGAP100F, Tsc1, and Mino. Together, this supports the functional distinctions for p.R47Q observed in the wing.

**Figure 4:**
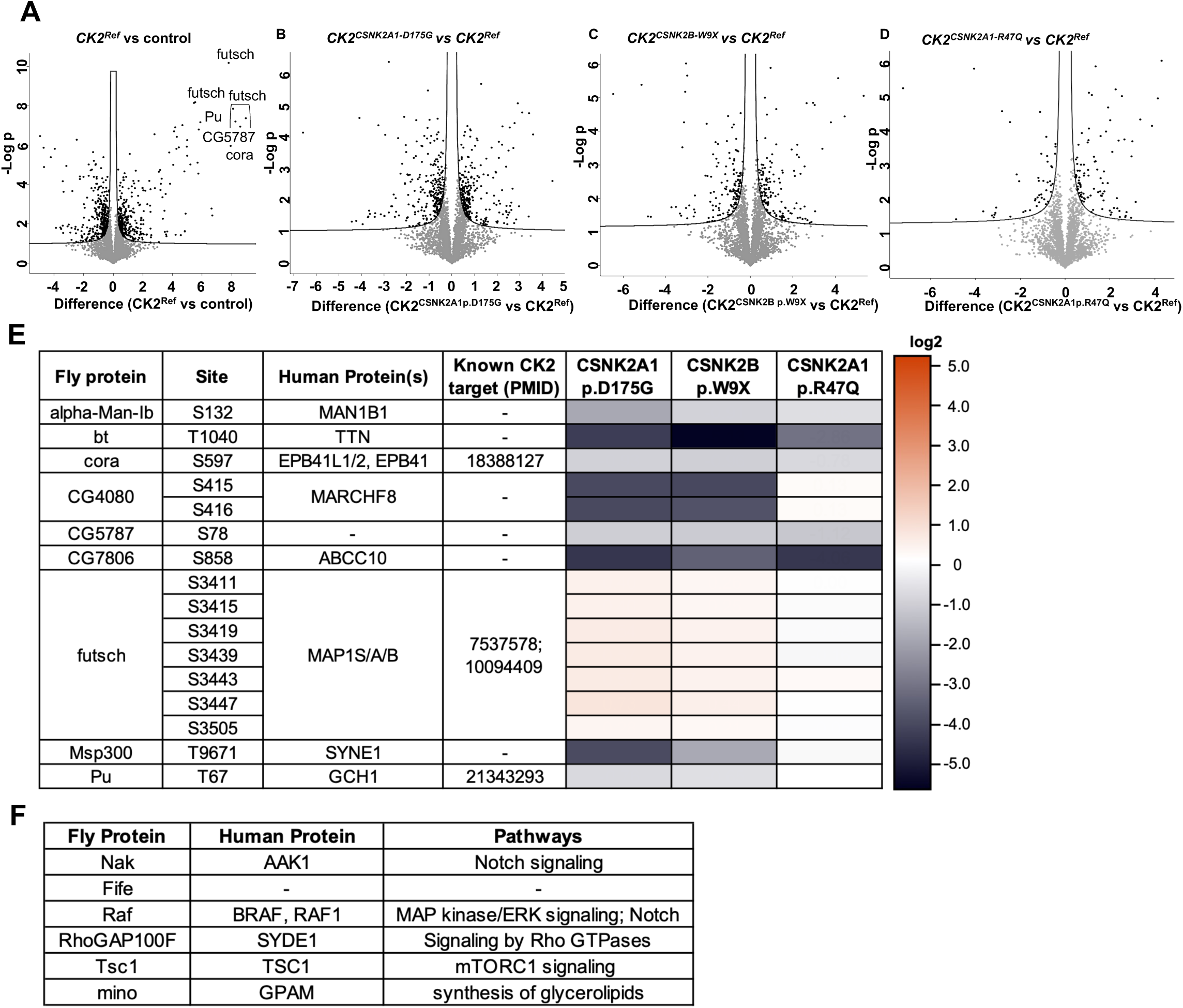
CK2 phospho-targets are decreased with LoF of variants and novel targets are identified with CSNK2A1 p.R47Q. (A-D) Volcano plots showing significantly different phosphorylation sites in fly heads expressing UAS constructs pan-neuronally with *nSyb-GAL4*, comparing control (A), *CK2^CSNK2A1^ ^p.D175G^* (B), *CK2^CSNK2B^ ^p.W9X^* (C), and *CK2^CSNK2A1^ ^p.R47Q^* (D) to *CK2^REF^*. (E) Heat map depicting top phospho-site changes from *CK2^REF^* (top right of A) in LoF variants *CK2^CSNK2A1^ ^p.D175G^* and *CK2^CSNK2B^ ^p.W9X^* and GoF variant *CK2^CSNK2A1^ ^p.R47Q^*. (F) Identified significant and specific *CK2^CSNK2A1^ ^p.R47Q^* phosphorylated proteins compared to *CK2^REF^*.

### Functional data of OCNDS and POBINDS variants in flies partially map to pathogenicity predictions and clinical severity

We next assessed if functional data for OCNDS and POBINDS pathogenic variants found in flies correlate with *in silico* pathogenicity prediction and reported clinical severity. OCNDS-related *CSNK2A1* variants, including their locations, *in silico* pathogenicity predictions, gnomAD allele frequencies, functional data in flies, and aggregated clinical severity scores are summarized in **Table 1**. *In silico* tools (aggregated REVEL and MetaLR scores) classified most variants as deleterious or uncertain. Notably, the recurrent p.K198R ‘hot spot’ variant was observed in two heterozygous individuals in gnomAD, whereas the remaining variants were absent from population databases. Functional studies predominantly supported LoF for p.Y50C, p.D175G, and p.R312W, which were all deemed deleterious by *in silico* scores. The four other variants showed functional evidence of partial LoF or mixed LoF/neomorphic behaviour (p.R47Q), which correlated with uncertain prediction scores.

**Table 1:**
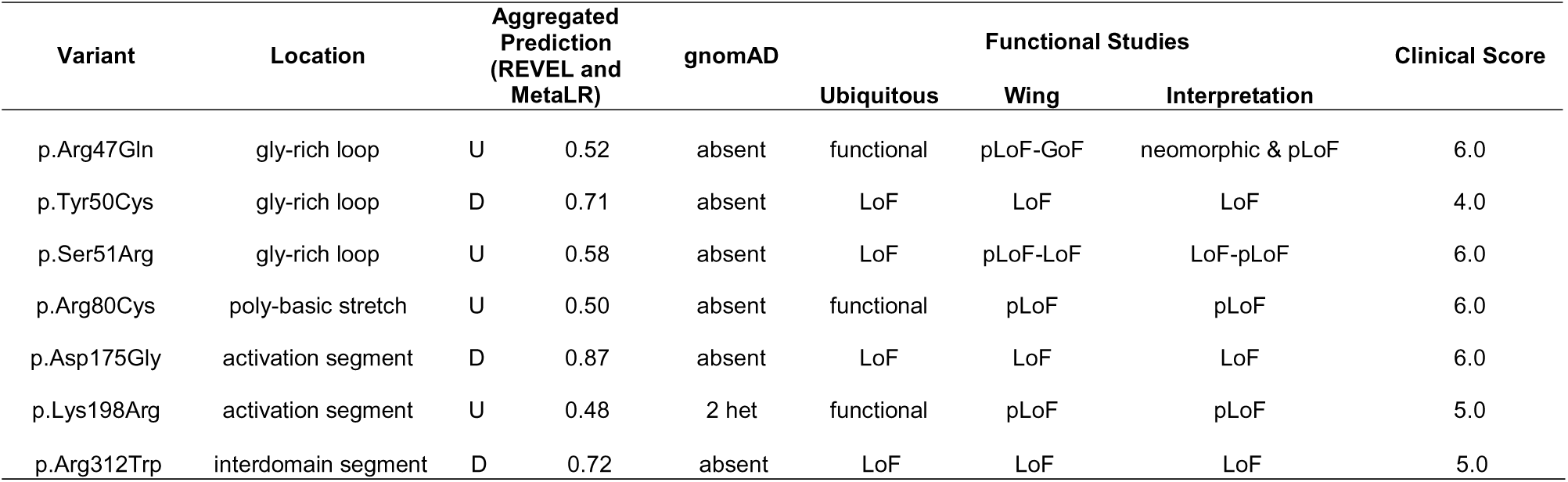
Summary of clinical scoring, *in silico* predictions, and functional data for OCNDS variants. D = deleterious, U = uncertain.

For POBINDS-related *CSNK2B* variants, the pathogenicity prediction, fly functional data, and clinical severity scores are summarized in **Table 2**. *In silico* predictions classified most variants as deleterious, with the p.H165R variant classified as uncertain. Functional studies largely supported a range of LoF effects (p.W9X, p.D32N, p.R86C, p.R111P, p.C137R and p.H165R). Clinical severity scores ranged widely from 2.6 (p.R111P) to 7.5 (p.H165R), highlighting marked phenotypic heterogeneity even among variants located in similar functional domains.

**Table 2:** Summary of clinical scoring, *in silico* predictions, and functional data for POBINDS variants. D = deleterious, U = uncertain.

| Variant | Location | Aggregated Prediction (REVEL and MetaLR) |  | gnomAD | Functional Studies |  |  | Clinical Score |
| --- | --- | --- | --- | --- | --- | --- | --- | --- |
|  |  |  |  |  | Ubiquitous | Wing | Interpretation |  |
| p.Trp9Ter | early beta | NA | NA | absent | LoF | LoF | LoF | 5.3 |
| p.Asp32Asn | early beta | D | 0.71 | absent | functional | LoF | LoF-pLoF | 4.7 |
| p.Arg86Cys | acidic loop - metal bdg | D | 0.88 | absent | functional | pLoF-LoF | LoF-pLoF | 4.0 |
| p.Arg111Pro | metal bdg site | D | 0.99 | absent | functional | pLoF | pLoF | 2.6 |
| p.Cys137Arg | metal bdg site | D | 0.87 | absent | LoF | LoF | LoF | 4.0 |
| p.His165Arg | interface to ck2a chain 1 | U | 0.67 | absent | pLoF | pLoF-LoF | LoF-pLoF | 7.5 |

### Constitutive and adult-specific neuronal knockdown of *CkII* reveals critical roles in development and behaviour

*Drosophila* CkIIα and CkIIβ are both 88% identical and 94/95% similar to human CK2α and CK2β at the amino acid level. All 12 missense variants analyzed above are conserved at the amino acid residue in the respective CkII proteins. Classical null alleles for *Drosophila CkIIα* and *CkIIβ* are homozygous lethal (26) and haploinsufficiency phenotypes are rare in *Drosophila* (27). Given that most pathogenic variants in OCNDS and POBINDS act as LoF, we aimed to model *CkIIα* and *CkIIβ* reduction in the nervous system using RNAi-mediated knockdown. Pan-neuronal or pan-glial knockdown of either *CkIIα* or *CkIIβ* using *nSyb-GAL4* or *repo-GAL4* causes lethality (**Fig. 5A**). *UAS-luciferase-RNAi* was used as a control, which results in viable flies. Since both *nSyb-GAL4* and *repo-GAL4* are expressed in embryonic and larval stages, these data suggest that CkII plays a critical role in both neurons and glia for organismal development.

**Figure 5:**
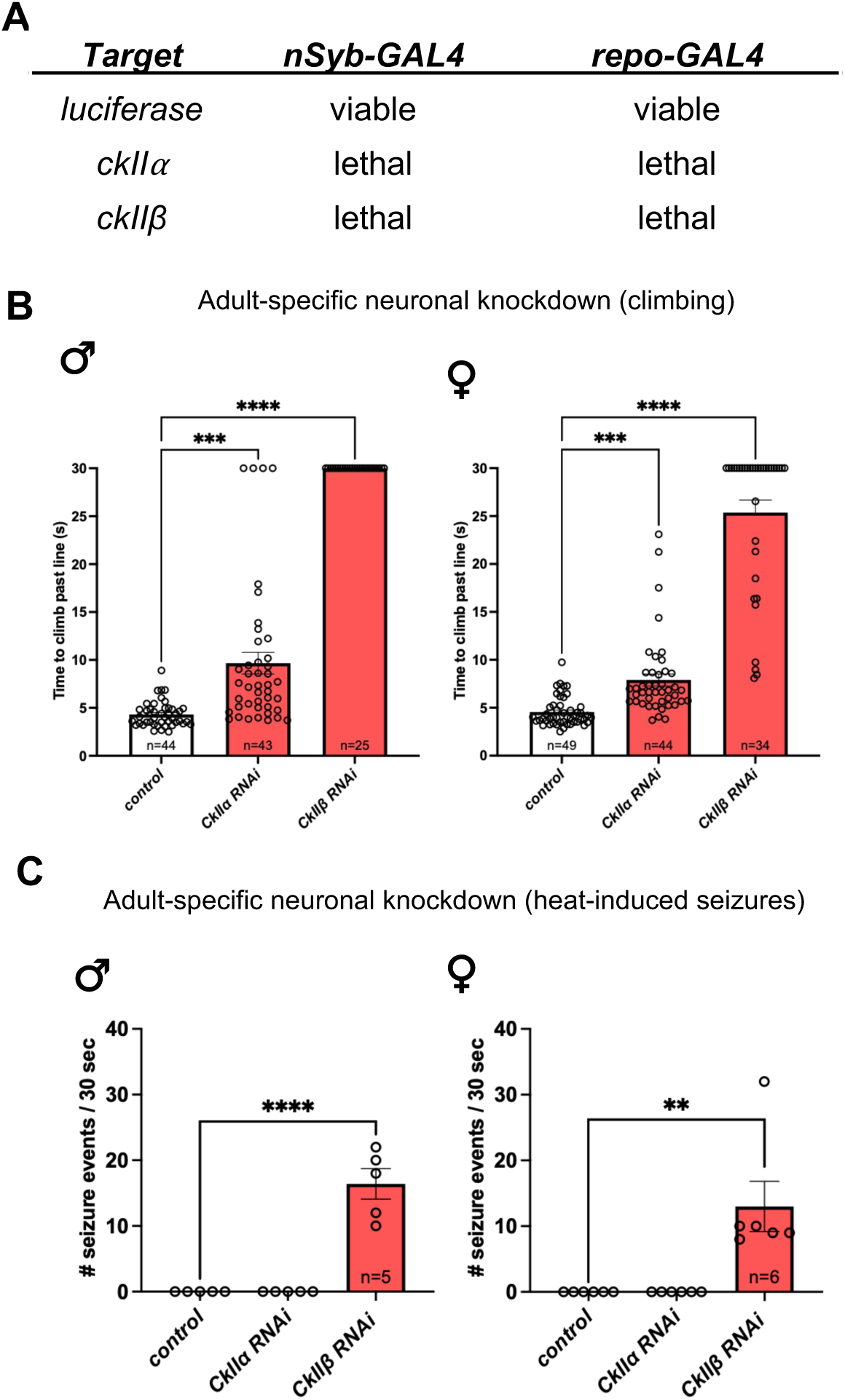
Neuronal and glial *CkII* is critical for development and adult-specific neuronal RNAi-mediated knockdown causes motor deficits and heat-induced seizures. (A) Table showing the results of RNAi-mediated knockdown of *CkII* in developing neurons and glia using *nSyb-GAL4* and *repo-GAL4*, respectively. Results were the same at 25°C and 29°C. (B & C) Quantification of negative geotaxis (climbing) and heat-induced seizures of 5-day-old adult flies upon pan-neuronal knockdown of *CkIIα* and *CkIIβ* using *elav^GS^ - GAL4* at 29°C in males and females. (B) Kruskal-Wallis test followed by Dunn’s multiple comparisons test, ***p<0.001, ****p<0.0001. (C) ANOVA followed by Tukey’s post hoc test, **p<0.01, ****p<0.0001. Climbing data were derived from a minimum of 15 flies per sex per genotype. For heat-induced seizure assay, data were derived from a minimum of five separate genetic crosses, with a minimum of 5 flies analyzed in each trial.

To avoid developmental lethality and examine the role of CkII in the adult nervous system, we employed adult-specific knockdown using the RU486 drug-induced GeneSwitch modified GAL4 system (28). Pan-neuronal, adult-specific knockdown of either *CkIIα* or *CkIIβ* using the *elav^GS^-GAL4* causes a significant increase in climbing time upon conducting a negative geotaxis climbing assay in both male and female flies at five days post-eclosion at 29°C (**Fig. 5B**). Adult-specific, pan-glial knockdown of either *CkIIα* or *CkIIβ* did not cause any significant climbing deficits (**Supplementary Material, Fig. S3**). Together, these data suggest that *CkII* has an important role in mediating motor behaviour in the mature nervous system.

To further model neural dysfunction, we also determined if adult-specific *CkII* knockdown resulted in any heat-induced seizure phenotypes. No seizure-like activity was observed in controls, or upon adult-specific neuronal knockdown of *CkIIα* using *elav^GS^ -GAL4* in either sex at five days post-eclosion at 29°C (**Fig. 5C**). However, adult-specific neuronal knockdown of *CkIIβ* displayed significant seizure-like behaviour. These data suggest that the β-subunit may have a critical role in mediating heat-induced seizure-like behaviour in flies.

### Pharmacologically enhancing Wnt/**β**-catenin partially rescues adult-specific CkII-related neuronal phenotypes

Acknowledging CK2’s broad functional landscape (29,30), we focused on enhancing the Wnt/β-catenin pathway as CK2 can potentiate this cascade at three levels via phosphorylation of: dishevelled - sequestering the destruction complex (31), β-catenin - increasing its resistance to degradation (32), and LEF-1 directly (33). Additionally, Wnt/β-catenin signalling has critical roles in neurodevelopment (34–37). Wnt agonists, Lithium Chloride (LiCl) and CHIR99021 enhance Wnt/β-catenin signalling via GSK3 inhibition (38). Treatment with either LiCl or CHIR99021 partially rescued climbing deficits observed upon adult-specific neuronal knockdown of *CkIIα* in male and female flies at 29°C (**Fig. 6A**). This suggests that Wnt/β-catenin signalling partially mediates motor phenotypes when *CkIIα* is reduced in the mature nervous system. For *CkIIβ*, treatment with LiCl or CHIR99021 partially rescued climbing deficits upon adult-specific pan-neuronal knockdown of *CkIIβ* in female but not in male flies at 29°C (**Fig. 6B**). However, Wnt agonist treatment partially rescued heat-induced seizure phenotypes observed upon adult-specific pan-neuronal knockdown of *CkIIβ* in both sexes (**Fig 6C**). These data suggest that Wnt/β-catenin partially mediates climbing and heat-induced phenotypes when *CkIIβ* is reduced in adult neurons, with potential sex differences in response depending on the phenotype.

**Figure 6:**
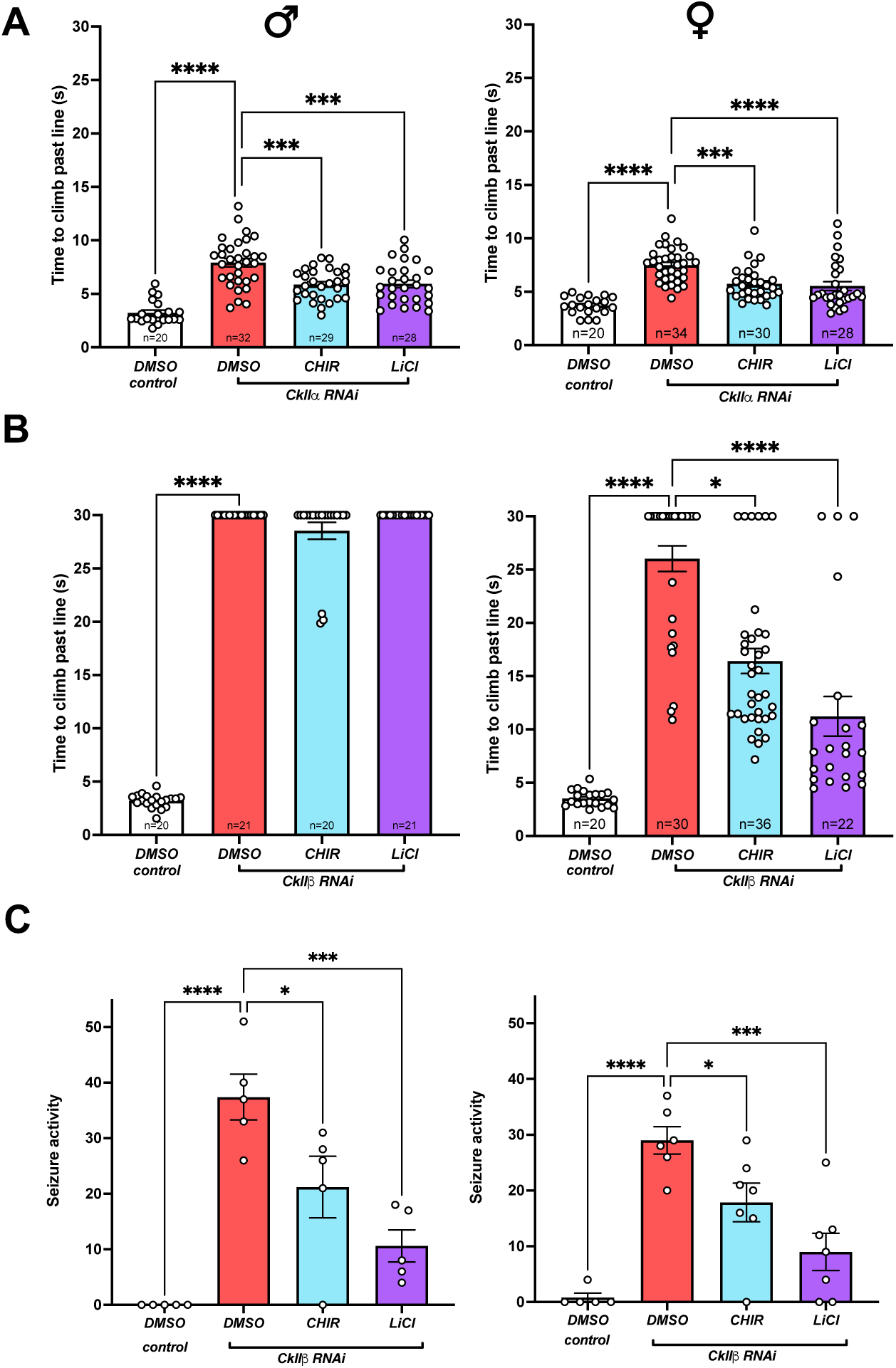
Drugs enhancing Wnt/β-catenin signalling partially rescue phenotypes associated with adult-specific pan-neuronal *CkII* reduction in flies. (A & B) Climbing activity upon adult-specific knockdown of *CkIIα* (A) or *CkIIβ* (B) in neurons. (C) Heat-induced seizure phenotype upon adult-specific knockdown of *CkIIβ* in neurons (C). CHIR = CHIR99021 at a final concentration of 200μM. LiCl at a final concentration of 50mM. (A & C) ANOVA and Tukey’s post hoc test, **p<0.01, ***p<0.001, ****p<0.0001. (B) Kruskal-Wallis test followed by Dunn’s multiple comparisons test, *p<0.05, ****p<0.0001. (A & B) Data were derived from a minimum of n=20 in each genotype. For heat-induced seizure assay, data were derived from a minimum of five separate genetic crosses, with a minimum of five flies analyzed in each trial.

## Discussion

We generated a transgenic toolkit in *Drosophila melanogaster* to examine human CK2 (*CSNK2A1* and *CSNK2B*) variants and enhance understanding of the molecular basis of CK2-related NDDs. A key experimental decision was to co-express *CSNK2A1* and *CSNK2B* in tandem - CK2^REF^. This approach was chosen because the majority of biochemical and structural studies indicate that CK2 functions optimally as a heterotetrameric holoenzyme (5,39–41). A few reports have suggested that individual subunits may retain partial activity when expressed alone (42–45), but our results clearly show enhanced functional consequences of overexpressing the human CK2 holoenzyme in flies compared with overexpression of cDNAs for the single subunits alone. However, single subunit (*CSNK2A1, CSNK2A2,* and *CSNK2B*) overexpression alone in the developing wing did show modest wing size reduction, indicating some function when overexpressed in this tissue. Regardless, this strategy highlights the importance of holoenzyme assembly of the human protein complex when expressed in *Drosophila*.

Functional assessment of seven OCNDS-related *CSNK2A1* pathogenic variants revealed that most (6/7 variants tested) act primarily through LoF or pLoF mechanisms. This aligns with independent *in vitro* kinase assays demonstrating reduced enzymatic activity for the majority of patient variants (11,46). In particular, the recurrent *hotspot* variant p.K198R (located in the activation segment) consistently showed pLoF behaviour. This variant is notable for its presence in two control individuals in gnomAD (47) and its pleiotropic clinical presentation, further illustrating the complex relationship between CK2 activity and phenotypic severity, even at the variant level (48). Intriguingly, protein analysis showed that the p.K198R about 50% the protein of CK2^REF^ for both subunits yet still was functional compared to variants expressed at levels closer to reference. For *CSNK2A1*, *in silico* pathogenicity predictions correlated with functional studies in flies where the deleterious scores mapped to p.Y50C, p.D175G, and p.R312W, which showed complete LoF in both ubiquitous and wing expression assays. The p.R312W showed near absent protein levels of CSNK2A1 which is aligned with expression studies in mammalian cells (11). Yet there were also severely reduced levels of CSNK2B. Both subunits are required for holoenzyme assembly, which increases stability, hence, the p.R312W change may decouple from CSNK2B lowering its stability. Moreover, CSNK2A1 (and CSNK2A2) contain all of the conserved consensus protein kinase motifs and possess constitutive kinase activity even in the absence of the regulatory CSNK2B subunit (49). However, association with CSNK2B modulates substrate selectivity, catalytic efficiency toward many substrates, and overall holoenzyme stability (5,7). The p.S51R, p.R80C, and p.K198R displayed pLoF and had scores in the uncertain range. The p.R47Q variant, showing neomorphic activity in the wing, also had an uncertain prediction score. All six pathogenic POBINDS variants showed a spectrum of pLoF to LoF, aligned with haploinsufficiency. The spectrum of functional impact for pathogenic *CSNK2B* was unexpected. This ranged from a mild pLoF for p.R111P to moderate pLoF variants, p.D32N, p.R86C, and p.H165R, and complete LoF variants, p.W9X and p.C137R. Fittingly, the p.C137R showed marked reduction of both subunits (although not statistically significant for CSNK2B). Intriguingly, the p.D32N showed near absent protein levels but still showed some partial function when expressed ubiquitously. *In silico* predictions were less reliable for variants affecting the regulatory subunit. Most were predicted to be deleterious, with only the p.H165R variant predicted as uncertain. This highlights the need for functional assessment. The discrepancy of *in silico* predictions between the catalytic and regulatory subunits has been noted before (50). Importantly, docking-site controlled UAS expression in flies revealed moderate variability of OCNDS and POBINDS variant protein expression by western blot. This suggests that variant-specific effects on stability and function need to be fully examined.

We were unable to establish a clear correlation between the variant impact observed in *Drosophila* and the aggregated clinical severity scores derived from published case reports for both *CSNK2A1* and *CSNK2B*. One intriguing standout was the mild pLoF of the p.R111P variant in *CSNK2B*, which had the lowest clinical score of the six POBINDS variants examined. However, this observed discrepancy likely stems from the static nature of individual case reports and inter-study variability in clinical phenotyping, particularly regarding the subjective descriptors used to characterize symptom severity.

We identified one particularly intriguing *CSNK2A1* variant, p.R47Q, located in the glycine-rich ATP-GTP-binding loop of CSNK2A1. Similar to *CK2^REF^*, the p.R47Q variant caused lethality upon ubiquitous overexpression, yet upon wing expression, it produced a clear novel phenotype (ectopic hairs), suggesting neomorphic/GoF activity. This was supported by neuronal expression studies identifying novel phospho-targets, including Nak, orthologous to human AAK1 (Adapter-associated Kinase 1). AAK1 has been shown to modulate Notch pathway activity (51). Altered Notch signalling at the dorsoventral boundary is a well-established cause of ectopic margin bristles in the *Drosophila* wing (52), which could explain the ectopic hairs observed in the wings. The Wnt/Planar Cell Polarity pathway could represents another putative pathway causing ectopic and disorganized hairs in the p.R47Q wing blade, especially as CK2 is known to potentiate Dishevelled (32). Five independent patients harbouring the p.R47Q variant have been reported in the literature (1,19,21,53,54). Intriguingly, two of these provided detailed case reports on the patients carrying the p.R47Q variant and described symptoms previously not associated with OCNDS, including clubfeet in both cases (53,54). Strikingly, one case reported an unusual pattern of scalp hair deemed difficult to comb (54). It should be noted that in our assays, the p.R47Q also exhibited pLoF. The data suggests that the p.R47Q harbours differential molecular consequences compared to other pathogenic *CSNK2A1* variants and it is tempting to speculate this may be responsible for this phenotypic expansion of the OCNDS spectrum. This highlights the value of functional testing using *Drosophila* to uncover variant-specific effects that may be missed by traditional LoF or *in vitro* approaches alone.

Although no other *in vivo* functional analysis of *CSNK2A1* or *CSNK2B* variants have been reported, a recent preprint expressed five OCNDS-associated *CSNK2A1* variants in zebrafish embryos, three of which overlapped with variants tested in our *Drosophila* studies (p.R47Q, p.Y50C, and p.K198R) (55). The authors observed variant-specific morphological defects and increased embryonic mortality, that could be rescued with co-injection of *CSNK2A1* reference. Although the mechanisms are not fully clear, this preprint suggested a mixture of LoF, dominant-negative, and potential GoF effects for pathogenic *CSNK2A1* variants. A clear distinction from our study is that only human *CSNK2A1* was overexpressed in zebrafish, whereas we co-expressed both subunits in *Drosophila*.

We next investigated the consequences of reducing endogenous *CkIIα* and *CkIIβ* in the *Drosophila* nervous system. Germline null mutant alleles for either *CkIIα* or *CkIIβ* are lethal (56–58). We found RNAi*-*mediated knockdown of either subunit during development in either neurons or glia results in pre-adult lethality. It has been previously shown that *CkII* knock down in neural stem cells with *insc-GAL4* leads to significant loss of neural stem cells in the late larval stage (59). We decided to focus on the role of CkII in the adult nervous system by inducing knock down after eclosion to highlight its role in the mature nervous system and assess neurobehavioural outcomes. We found that CkII in adult neurons is important for maintaining normal motor function and seizure susceptibility. We only observed heat-induced seizure-like behaviour upon *CkII*β knockdown in adult-neurons. This could be explained by the varying strengths of the *CkIIα* and *CkII*β UAS-RNAi lines and/or reflective of different expression levels of the two genes to begin with. Interestingly, seizures are much more common in POBINDS (∼90%) compared to OCNDS (∼30%) (60).

Although CK2 is known to phosphorylate over 300 substrates (29,30), we chose to focus on modulating the Wnt/β-catenin pathway for CK2 in potentiating this pathway at multiple levels (31–33) as well as its known roles neurodevelopment and genetic alterations in NDDs (35,61–65). Pharmacological enhancement of Wnt/β-catenin signalling with LiCl or the GSK3 inhibitor, CHIR99021, partially rescued the climbing and seizure phenotypes in adult-specific CkII knockdown flies. Patient-cells from POBINDS have shown reduced β-catenin phosphorylation and nuclear localization, and disruption of canonical Wnt signalling (66). We did not observe any rescue of climbing deficits in male flies upon *CkIIβ* knockdown in adult neurons. This may be due to the extreme motor phenotype observed in this group. Regardless, these results indicate that GSK3 inhibition can partially mediate the behavioural deficits observed, suggesting that Wnt pathway modulation may hold therapeutic potential for CK2-related disorders. However, issues of toxicity, blood-brain barrier penetration, and pediatric safety will require careful evaluation. Drug concentrations were based on prior studies; further optimization of additional concentrations and/or combination treatment with drugs targeting other CK2-affected pathways (highlighted from our phosphoproteomic screen) may yield more complete rescue.

The tandem co-expression of human *CSNK2A1* and *CSNK2B* in *Drosophila* provides a rapid approach for functional assessment. By comparing patient variants to the *CK2^REF^*, ubiquitous overexpression distinguishes strong LoF effects, while tissue-specific assays in the developing wing additionally reveals pLoF and GoF phenotypes. Considering the growing number of Variants of Uncertain Significance (VUS) in clinical databases (67), there is a need for functional testing to support diagnosis. Most VUS are missense changes, which is why we focused on pathogenic missense changes in *CSNK2A1* and *CSNK2B*. The *in vivo* readouts we have developed may help re-classify VUS and prioritize variants for further study. This would accelerate clinical interpretation in OCNDS and POBINDS, where functional data remain limited and diagnostic uncertainty is common. Differential variant function may be critical for developing distinct treatment strategies, with LoF variants potentially benefiting from CK2 gene replacement or Wnt enhancement, while GoF/neomorphic variants may also require inhibition or pathway-specific targeting.

The limitations of this study include the reliance on transgenic overexpression in flies that may not fully recapitulate endogenous expression levels, potentially amplifying or masking certain variant effects. Confirmation of key findings, such as the neomorphic/GoF activity of p.R47Q, in human cell lines or mammalian models would strengthen translational relevance. Moreover, expression of human CK2^REF^ was performed in a wild-type background, thus interactions with the endogenous fly CkII complex cannot be ruled out. Future studies could generate and examine endogenous knock-in flies in heterozygous and homozygous animals, if viable. In summary, we have developed a transgenic *Drosophila* toolkit to rapidly distinguish LoF, pLoF, and GoF effects of *CSNK2A1* or *CSNK2B* variants, highlighting variant-specific mechanisms in OCNDS and POBINDS. These insights may advance precision diagnosis for VUS and lay the groundwork for targeted therapies in CK2-related NDDs.

## Materials and Methods

### Fly husbandry and stocks

Fly stocks were maintained at ambient room temperature (22°C). Standard molasses-based fly food was used for the maintenance of the stocks. The following stocks were used in this study:

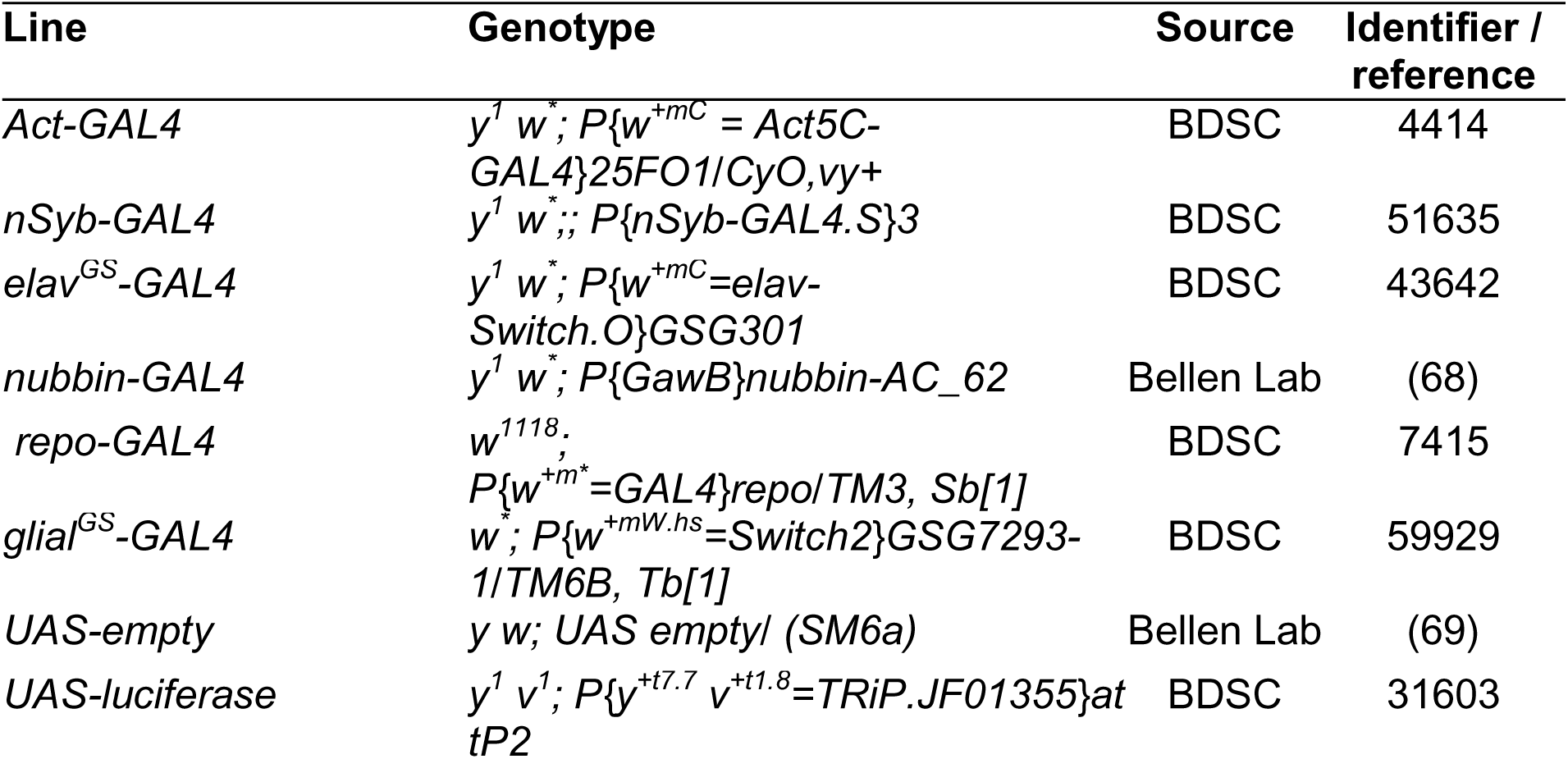
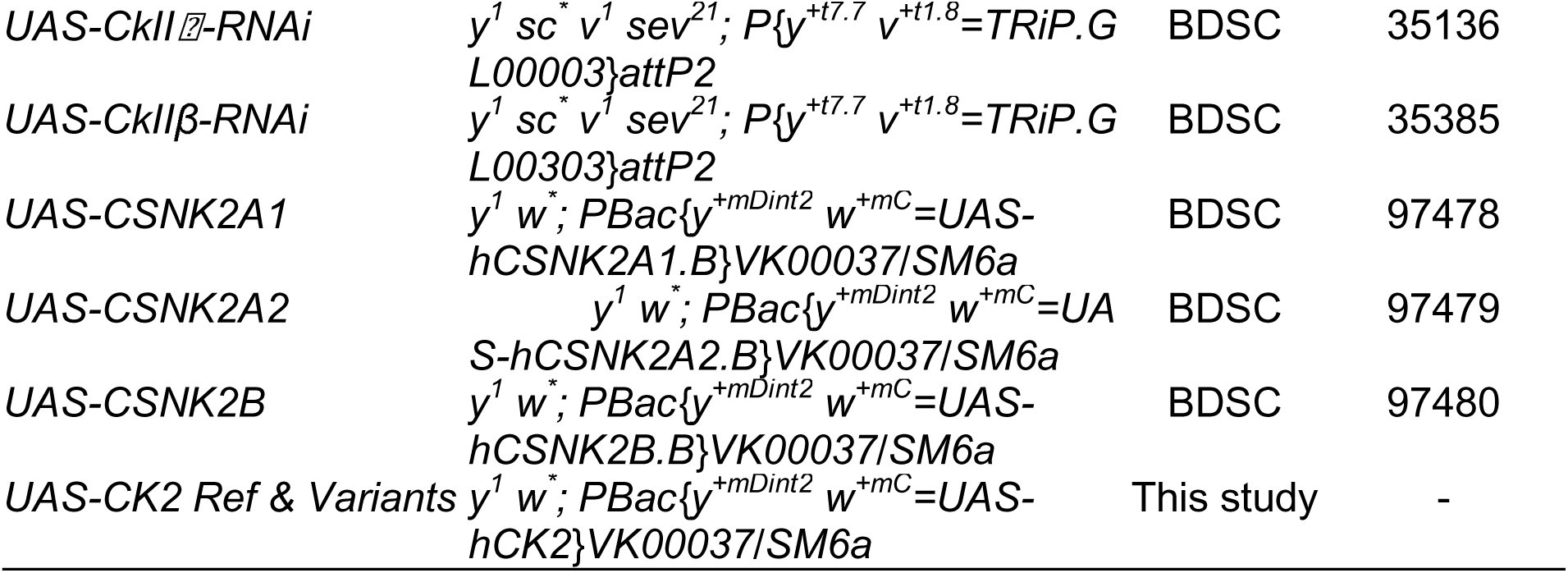

### Generation of *UAS-CSNK2A1::T2A::CSNK2B* transgenic flies

Briefly flies were generated as previously (70). Briefly, *UAS-CNK2A1::T2A::CSNK2B* cDNA was synthesized (Twist Bioscience) using NM_001895.3 and NM_001320.5 with a T2A sequence (GAGGGCCGCGGCAGCCTGCTGACCTGCGGCGATGTGGAG-GAGAACCCCGGGCCC) in between. Oligonucleotides used for SDM are as follows:

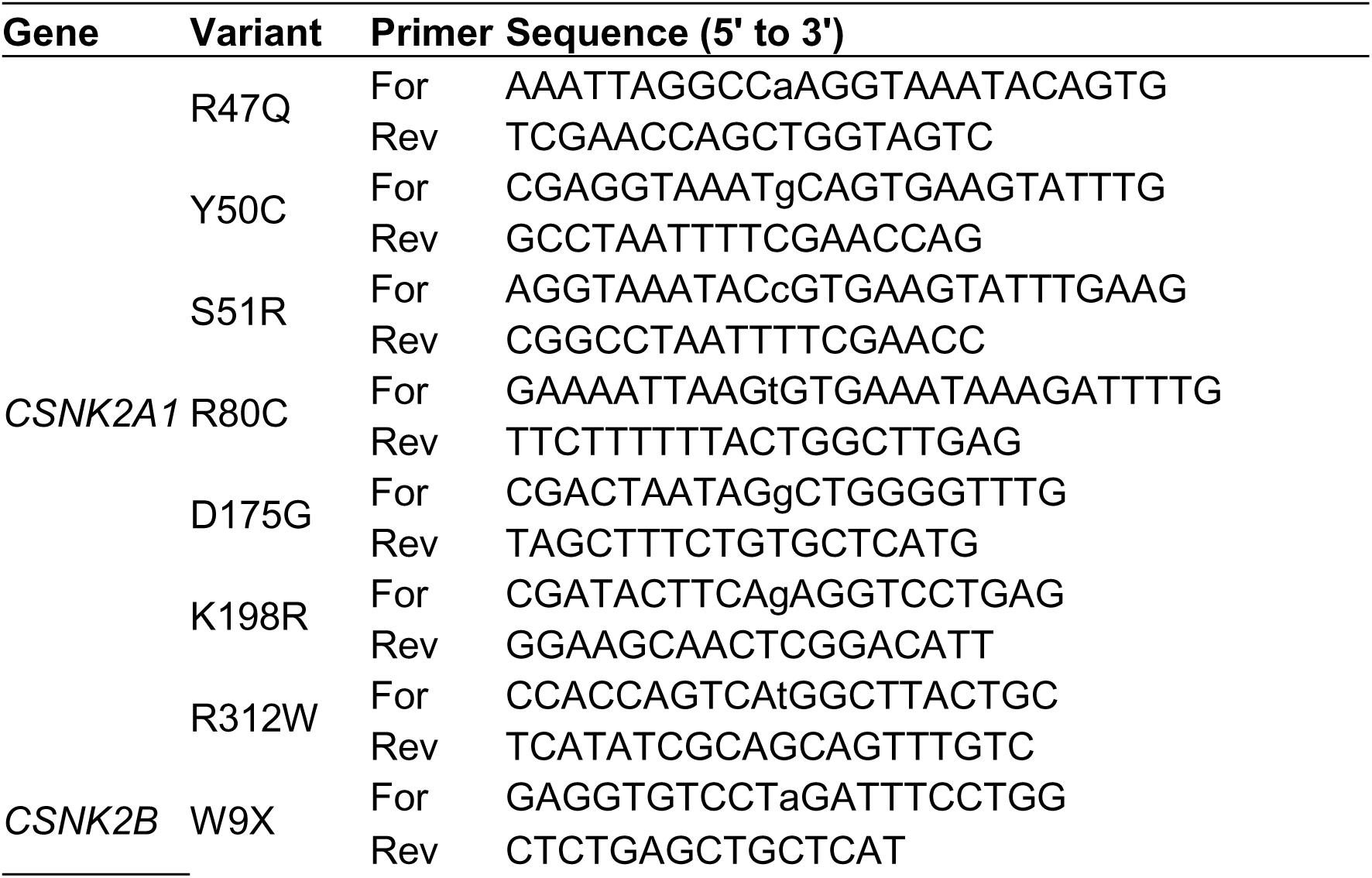
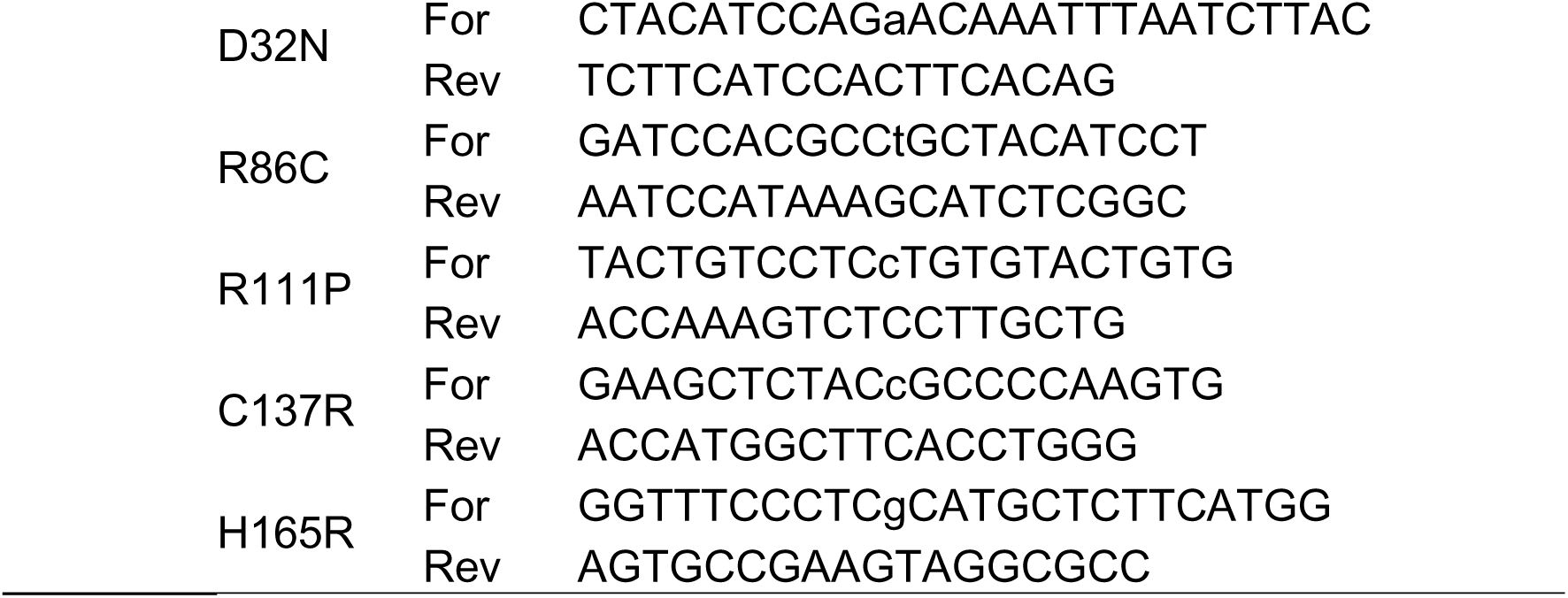

Pathogenic variants and reference cDNA for *CSNK2A1* (NM_001895.3) and *CSNK2B* (NM_001320.5) were made in Gateway-compatible DONR plasmids by site-directed mutagenesis (SDM) using the Q5 SDM Kit (New England Biolabs (NEB)). 10 ng of *CK2* reference DONR plasmid as template DNA was added to 12.5 μL of Q5 Hot Start High-Fidelity master mix, with 0.5μL forward and reverse mutagenesis primers, and nuclease-free water were mixed and underwent a polymerase chain reaction (PCR). After PCR, a kinase, ligase and DpnI enzymatic reaction was completed using 1μL of the PCR product, 5 μL of the 2XKLD reaction buffer (NEB), 1 μL of KLD enzyme mix (NEB) and 3 μL nuclease-free water, which was then added to a chemically competent DH5α cells and incubated on a plate with vector antimicrobial resistance for transformation. The transformed colonies were harvested for a Miniprep using the Invitrogen Kit (K210011) for DNA purification and extraction. The colonies were then sent to The Hospital for Sick Children (SickKids) for Sanger sequencing verification. Using an Invitrogen Kit, Gateway Cloning TM (ThermoFisher) was done to transform the DONR plasmids into to a fly compatible destination vector (pGW-HA.attB) (71). 150 ng of empty pGW vector, 150ng of DONR vector, 1 μl of LR Clonase II and tris-ethylenediamine tetra-acetic acid (TE) butter were incubated overnight at room temperature. 1 μL of Proteinase K was added the following day and was incubated in a 37°C water bath for 10 minutes to stop the reaction. 1 μL of this solution was added to competent cells for transformation and was plated onto agar plates with ampicillin. Using the Invitrogen Kit (K210004), Midiprep was completed for DNA extraction and purification. pGW plasmids containing the *CK2* reference or variants were sent to Plasmidsaurus for whole plasmid sequencing verification by next-generation sequencing. Purified DNA was injected into VK37 (PBac{y[+]-attP}VK00037) docking site (**72**) by ϕC31-mediated transgenesis (GenomeProlab). All *UAS-cDNA* were inserted into the same docking site, controlling for any positional effects.

## Drosophila viability

Both sexes were analyzed and viability was assessed as previously (15). Progeny were visually genotyped and counted. Flies possessing the desired genotype were identified by the absence of the dominant morphological marker from the parental balancer chromosome. Mendelian ratios were determined by first isolating and quantifying progeny lacking the marker. This count was divided by the total number of flies scored to obtain the proportion of observed progeny. The observed proportion was then normalized to the theoretical Mendelian expectation. Ratios were calculated independently for each sex and temperature.

## Drosophila wing analysis

Adult wing dissection, mounting, imaging and phenotypic quantification were performed as previously (**73**). Briefly, using the *nubbin-GAL4* driver, crosses were made to over-express human cDNA in the developing fly wings. 5-7 *nubbin-GAL4* virgin females were crossed with 3-5 male transgenic variant flies harbouring human cDNA. Crosses were completed in both 25°C and 29°C. 7-10 days after the cross was made, progeny were collected and kept in 70% ethanol. Adult flies were dissected for their wings and was mounted on microscope slides using wing mounting medium 3:1 dilution of CMCP-10 mounting medium (Masters, Wood Dale, IL) and lactic acid. A coverslip was added to protect the wings, and the borders were sealed with clear nail polish to prevent cover slips from sliding. 20 wings per sex, genotype and temperature were mounted on each slide. Slides were imaged using the Axioscan 7 by Zeiss. Both sexes were examined for each genotype and temperature. Wings were examined for abnormal morphological phenotypes including the size of the wing, vein defects, bristle loss, blisters and hair whirls. Wing size was determined by measuring the surface area of each wing using ImageJ by Fiji, which were then normalized to the wing sizes of the Lac z controls. The presence of blisters was counted prior to wing dissection to avoid miscounts after being flattened and mounted. Lastly, the presence of ectopic hair phenotypes were recorded.

## Protein analysis

Protein analysis was performed as previously described (**70**). Briefly, 5-day-old flies were separated depending on their sex and 5 heads were isolated and collected in cold RIPA buffer 30 μL with Halt^TM^ protease inhibitor (ThermoFisher #1862209). The fly heads were then lysed using manual pestles (3X for 10 seconds each rep on ice). Samples were then centrifuged for 1 min at 12000 RPM. Tissue lysate with Laemmli Buffer was boiled for 5 minutes and resolved by SDS-PAGE followed by transferring the proteins onto a PVDF membrane. Primary antibodies used were anti-CSNK2A1 (rabbit, ThermoFisher P5-28686), anti-CSNK2B (rabbit, ThermoFisher P5-27416), and anti-Actin (rabbit, Abcam AB227387). Secondaries were goat anti-rabbit HRP (ThermoFisher 31460).

## Phospho-proteomics

To obtain global proteomics from flies, 4% SDS in 100 mM TRIS, pH 8.5 used to mechanically dounce five fly heads in 30 μL of lysis buffer. Samples were vortexed for 10 min, and then centrifuged for 10 min at 15000g. Bradford assay was done using Pierce™ Detergent Compatible Bradford Assay Kit (ThermoScientific, #23246) as per manufacturer’s protocol to determine protein concentration. 30 μg from each sample was transferred to a new vial and had their volumes normalized using 4% SDS in 100 mM TRIS, pH 8.5. Samples were reduced, alkylated, and quenched the samples under the following conditions: Reduction: 10 mM DTT final and incubation at 57°C for 30 min; Alkylation: 50 mM IAA final and incubation at room temp in the dark for 45 min; and Quench: 17 mM DTT final and incubation at room temperature for 10min. Samples were confirmed to be pH∼8.0 before the samples underwent a single-pot solid-phase-enhanced sample preparation (SP3) protocol: Firslty, prior to SP3 treatment, two types of carboxylate-modified SeraMag Speed beads (Cytiva, cat no. 45152105050250 and 65152105050250) were combined in a ratio of 1:1 (v/v), rinsed, and reconstituted in water at a concentration of 20 μg solids/ μL. The reconstituted bead solution was added to the samples for a final concentration of 0.37 μg/μl. To promote proteins binding to the beads, acetonitrile was added to a final concentration of 70% (v/v) and samples were incubated at room temperature on a tube rotator for 18 min. Subsequently, beads were immobilized on a magnetic rack for 1 min. The supernatant was discarded and the pellet was rinsed 2X with 200 μl of 70% ethanol and 1X with 180 μl of 100% acetonitrile while on the magnetic rack. Rinsed beads were resuspended in 50 mM HEPES, pH 8.0 supplemented with trypsin (Promega, cat no. V5111) at an enzyme-to-protein ratio of 1:25 (w/w) and incubated for 16 hours at 37°C. The tubes were immobilized on a magnetic rack once again and the supernatant was collected in a clean tube. To ensure greater recovery, the beads were removed from the rack and sonicated in a bath sonicator with water and collected in their respective tubes from the previous step. The Pierce Quantitative Fluorometric Peptide Assay (Fisher Scientific, cat no. 23290) was used to determine the peptide concentration. An aliquot from each sample was taken to acidify to pH2-3 using formic acid. 200 ng from the acidified aliquot of each sample was loaded into Evotip Pure (Evosep, cat no. EV2011) using manufacturer protocol for MS submission and analysis.

For phospho-proteomics, 10μg aliquot from each sample digest was taken and had an equal volume of 2x binding buffer (80% acetonitrile and 10% trifluoroacetic acid in 0.2M glycolic acid) added to it. In a separate vial, a 1:1 mixture of Ti-IMAC HP (Magresyn, cat no. MR-THP010) and Zr-IMAC HP (Magresyn, cat no MR-ZHP010) beads were added for a 1:20 (w/w) ratio of peptide to combined bead mixture. After aliquoting, the bead vials had 80 μl of 1x binding buffer (80% acetonitrile and 5% trifluoroacetic acid in 0.1M glycolic acid) added to them. The beads were then immobilized on a magnetic rack and had the supernatant removed. The beads were washed further by adding another 120μl of 1x binding buffer, pipette mixing, immobilizing the beads on a magnetic rack, and removing the supernatant. Each peptide sample was transferred into their own vial containing washed beads. To bind the phosphopeptides onto the beads, the samples were vortexed for 20 min at 800 rpm. At the end of incubation, the supernatant was removed and discarded. 150 μl of 1x binding buffer was added to each sample and pipette mixed. After all samples had gotten mixed, they were placed back on the magnetic rack to immobilize the beads and had their supernatant removed. The vials were taken off the magnetic rack afterwards. Next, 200 μl of wash buffer 2 (1% trifluoroacetic acid in 80% acetonitrile) was added to each sample and pipette mixed. After all samples were mixed, they were placed back on the magnetic rack to immobilize the beads and had their supernatant removed. The vials were taken off the magnetic rack afterwards. 200μl of wash buffer 3 (0.2% trifluoroacetic acid in 10% acetonitrile) was added to each sample and pipette mixed, then placed back on the magnetic rack as above. To elute the phosphopeptides off the beads, 90 μl of 2% NH4OH was added to each vial and vortexed for 1 min at 800rpm. Vials were placed on a magnetic rack to immobilize the beads and transfer the supernatant to a vial containing 20 μl of 10% formic acid to acidify the samples to pH∼2.0-3.0. The entire phosphopeptide enriched sample was loaded into Evotip Pure (Evosep, cat no. EV2011) using manufacturer protocol for MS submission and analysis.

Enriched phosphopeptide samples were analyzed by data-dependent acquisition (DDA) LC–MS/MS on a Bruker timsTOF Pro 2. Raw Bruker .d files were processed using FragPipe v23.2 with MSFragger v4.4.1, MSBooster v1.3.30, Percolator v3.7.1, PTMProphet v6.3.2, Philosopher v5.1.3-RC7, and IonQuant v1.11.20. Spectra were searched against a *Drosophila melanogaster* UniProt proteome database supplemented with contaminants and reversed decoys (Uniprot proteome UP000000803 downloaded January 13, 2026, 13,824 target fruit fly sequences). Trypsin was specified as enzyme allowing two missed cleavages, peptide length was set to 7–40 amino acids. Precursor and fragment mass tolerances were set to 20 ppm. Carbamidomethylation of cysteine was specified as a fixed modification. Oxidation of methionine, phosphorylation of serine/threonine/tyrosine, and deamidation of asparagine/glutamine were specified as variable modifications, with up to four variable modifications per peptide. Phosphorylation-site localization was performed with PTMProphet, and site-level quantification was performed using IonQuant with MS1-based label-free quantification, normalization, requantification, and matching between runs enabled. Feature extraction used 10 ppm m/z tolerance, 0.4 min RT tolerance, 0.05 ion-mobility tolerance, at least two isotopes, at least five scans across the peak, and at least two ions. Matching between runs used 0.7 min RT tolerance, 0.05 ion-mobility tolerance, up to 10 donor runs, and no minimum correlation threshold. Phosphosite-level reports were generated using a minimum localization probability of 0.75.

In total, 15,591 phosphorylation sites were quantified. Downstream analysis was performed in Perseus after log2 transformation of site intensities. For each pairwise comparison, only sites quantified in all biological replicates of at least one of the two compared conditions were retained. Differential phosphorylation was assessed in Perseus using its default multiple-testing correction procedure, with significance defined at 5% FDR. Volcano plots and principal component analysis were generated in Perseus. Sites quantified in all replicates of one condition and absent from all replicates of the compared condition were annotated as condition-specific when they did not pass the statistical test, for example because of high replicate variability.

For module-level analysis, phosphosite quantification data were exported from FragPipe/IonQuant in Perseus-style format, yielding 12,431 sites across five conditions with four biological replicates each: UAS_empty, CK2_ref, CK2_W9X, CK2_D175G, and CK2_R47Q. Complete-case sites quantified in all 20 samples were retained for omnibus testing, resulting in 6,148 sites. These sites were tested by one-way ANOVA across the five conditions using replicate-level measurements. Statistical significance was controlled by permutation-based FDR, using 1,000 permutations of condition labels while preserving group sizes. Sites with q_permFDR ≤ 0.05 were retained as condition-responsive, yielding 278 sites. These sites were z-scored within each site and clustered by their five-condition mean phosphorylation profiles into five response modules. Module activity was calculated for each individual sample as the mean z-scored phosphorylation signal across all sites assigned to the module.

## Drosophila lifespan and behaviour

### Climbing Assay

Climbing assays (also known as negative geotaxis) was performed as previously (73). 24 hours before testing, flies were put on standard fly food individually. At the time of testing, individual flies were transferred to a sterile vial lacking food and habituated for 1-2 minutes. Flies were tapped to the bottom of the vial and assessed for negative geotaxis, given 30 s to reach the 7-cm mark on the vial.

### Heat-induced seizure assay

Heat-induced seizure assay was performed as previously (74). 5 days after the flies eclosed, they were put on appropriate fly foods. The flies were then transferred to a clean vial lacking food and were habituated 5-15 min before the vial was submerged in a 42°C water bath for 30s. Seizure activities were defined as failure to maintain an upright posture combined with wing and body twitching. The number of seizure events in the duration of 30s was counted and scored for each group.

### Drug treatment in flies

Adult-specific knockdown studies using GeneSwitch GAL4 drivers was performed as previously (**75**). For the *elav^GS^-GAL4* driver, Mifrepristone from Sigma Aldrich (M8046, 500mg) was diluted with 100% ethanol (15.5 mL) to the desired concentration of 75 mM. 10μL of the diluted solution is added to 2.5 mL of standard molasses-based fly food. New food was provided every three days. Lithium Chloride from ThermoFisher Scientific (L121500) was diluted to 12.5 M in distilled water. 5 μL of this solution was mixed into 2.5mL of standard molasses-based fly food to make a final concentration of 50 mM. CHIR99021 from MilliporeSigma Canada Ltd (SML1046-25MG) was diluted to a 100 mM stock solution in DMSO. 5 μL of this solution was mixed with 2.5 mL of standard molasses-based fly food to make a final concentration of 200 μM. Control treatment was DMSO. New drug food was given every three days.

### Clinical scoring

Pubmed and Google Scholar databases were accessed to review articles containing clinical information on the seven selected *CSNK2A1* variants and the 6 selected *CSNK2B* variants. Search terms included “CK2A” or “Okur Chung Neurodevelopmental Syndrome (OCNDS)” AND “variants” for CK2A-related syndrome, and “CK2B” OR “Poirier Bienvenu Neurodevelopmental Syndrome (POBINDS)” AND “variants” for CK2B-related syndromes. Clinical information, including sex, age, and the presence of characteristics (DD/ID, seizure, microcephaly, dysmorphism and congenital anomaly, etc.), was collected from the selected articles, and a clinical table was developed. The clinical score was divided into three independent scores of neurodevelopmental, neurological and non-neurological scores. The neurodevelopmental score for OCNDS or *CSNK2A1-*related syndrome and POBINDS or *CSNK2B*-related syndrome was determined by the severity of developmental delay and intellectual disability. Scores were divided into four categories, including one point for mild ID and up to four points for profound ID. The neurological score for OCNDS or *CSNK2A1*-related syndrome had a maximum of two points. The categories included absence of seizures, scoring 0, treatable seizures scoring 1, and untreatable seizures with a score of 2. The other two points came from the presence of microcephaly, two points for the presence and zero point for absence of microcephaly. For POBINDS or *CSNK2B*-related syndrome, the neurological score had a maximum of two points for seizure with treatment and zero points for seizure without treatment (all individuals with POBINDS had seizures). The systemic score for OCNDS or CSNK2A1-related syndrome included the presence of facial dysmorphism, with a maximum score of 1 point for the presence of facial dysmorphisms. The systemic score for POBINDS or *CSNK2B*-related syndrome included the presence of facial dysmorphism and congenital anomaly, with a maximum score of 2 points for the presence of each characteristic. The total clinical score was calculated by combining the three scores together. The maximum clinical score for individuals with OCNDS was 9. For individuals with POBINDS, the maximum clinical score was 8.

### *In-silico* predictions

*In-silico* predictors were analyzed using the Franklin Genoox platform (https://franklin.genoox.com). Aggregated scores were derived from including REVEL (76) and MetaLR (77).

## Acknowledgements

We would like to thank Drs. Eric Hall and David Litchfield for critical comments on the project/manuscript. We thank Catherine Bland and Adedayo Oladejo for technical assistance during the project.

## Author contributions

Yina Her (Formal analysis, Investigation, Methodology, Visualization, Writing—original draft, Writing—review & editing), Danielle M. Pascual (Investigation), Ying Lao (Investigation, Methodology), Harsimran Kaur (Investigation), Alondra Griffiths (Investigation), Robert Beattie (Resources), Bradley W. Doble (Resources), Patrick Frosk (Formal analysis), Rene Zahedi (Formal analysis, Investigation), Paul C. Marcogliese (Conceptualization, Funding acquisition, Formal analysis, Investigation, Methodology, Visualization, Resources, Writing—original draft, Writing—review & editing).

## Supplementary Legends to Figures

**Supplementary Figure 1:** Wing pouch expression of pathogenic *CSNK2A1* or *CSNK2B* variants reveals a spectrum of LoF with the CSNK2A1 p.R47Q acting as a GoF. Representative wing images upon wing-specific expression of *UAS-CSNK2A1* variants (A) and *UAS-CSNK2B* (E) using *nub-GAL4* at 29°C. Quantification of wing size and presence of blisters for *UAS-CSNK2A1* (B & D), *UAS-CSNK2B* (F & G). (C) 10X zoom image of the CSNK2A1 p.R47Q variant showing ectopic hair phenotype. (B & F) ANOVA followed by Tukey’s post hoc test, ****p<0.001. (D & G) Kruskal-Wallis test followed by Dunn’s multiple comparisons test, was completed with significance defined as *p<0.05, **p<0.01, ***p<0.001, ****p<0.001. Data were derived from three independent genetic crosses, with a minimum of n=20 in each cross. All data derived from female flies.

**Supplementary Figure 2:** Wing pouch expression of *CSNK2A1* or *CSNK2B* NDD-associated variants reveals loss of bristle phenotypes compared to reference. Quantification of bristle loss upon wing-specific expression of *CSNK2A1* variants (A & B) and *CSNK2B* (C & D) using *nub-GAL4* at 29°C. Kruskal-Wallis test followed by Dunn’s multiple comparisons test, was completed with significance defined as ****p<0.001. Data were derived from three independent genetic crosses, with a minimum of n=20 in each cross. Sex indicated by Greek symbols.

**Supplementary Figure 3:** Adult-specific glial knockdown of *CkII* does not significantly alter climbing behaviour. Climbing activity upon pan-glial knockdown of *CkIIα* and *CkIIβ* using the *glial^GS^ -GAL4* driver at 29°C in 5-day-old flies. ANOVA and Tukey’s post hoc test. Data were derived from a minimum of n=30 in each genotype.

## Abbreviations

AAK1: Adapter-associated Kinase 1
CK2: Casein Kinase 2
CK2α: Casein Kinase 2 alpha subunit
CK2β: Casein Kinase 2 beta subunit
GoF: Gain-of-Function
GS: GeneSwitch
LiCl: Lithium Chloride
LoF: Loss-of-Function
OCNDS: Okur-Chung Neurodevelopmental Syndrome
NDDs: Neurodevelopmental disorders
nSyb: neuronal synaptobrevin
pLoF: partial Loss-of-Function
POBINDS: Poirier-Bienvenu Neurodevelopmental Syndrome
RNAi: RNA interference
UAS: Upstream Activation Sequence
VUS: Variant of Uncertain Significance

## References

1. Okur, V., Cho, M.T., Henderson, L., Retterer, K., Schneider, M., Sattler, S., Niyazov, D., Azage, M., Smith, S., Picker, J., et al. (2016) De novo mutations in CSNK2A1 are associated with neurodevelopmental abnormalities and dysmorphic features. Hum Genet, 135, 699–705.

2. Poirier, K., Hubert, L., Viot, G., Rio, M., Billuart, P., Besmond, C. and Bienvenu, T. (2017) CSNK2B splice site mutations in patients cause intellectual disability with or without myoclonic epilepsy. Hum Mutat, 38, 932–941.

3. Li, D., Zhou, B., Tian, X., Chen, X., Wang, Y., Hao, S., Zhang, C. and Hui, L. (2024) Genetic analysis and literature review of a Poirier–Bienvenu neurodevelopmental syndrome family line caused by a de novo frameshift variant in CSNK2B. Mol. Genet. Genom. Med., 12, e2327.

4. Li, X., Wang, S., Liu, X., Wang, Z., Lv, N., Wang, S. and Yang, W. (2025) A Case of CSNK2A1 Gene Variant Causing Okur-Chung Syndrome and Analysis of the Clinical Phenotypic Spectrum. Mol. Genet. Genom. Med., 13, e70166.

5. Borgo, C., D’Amore, C., Sarno, S., Salvi, M. and Ruzzene, M. (2021) Protein kinase CK2: a potential therapeutic target for diverse human diseases. Signal Transduct Target Ther, 6, 183.

6. Pinna, L.A. (2002) Protein kinase CK2: a challenge to canons. J Cell Sci, 115, 3873–3878.

7. LITCHFIELD, D.W. (2003) Protein kinase CK2: structure, regulation and role in cellular decisions of life and death. Biochem. J., 369, 1–15.

8. Turowec, J.P., Duncan, J.S., French, A.C., Gyenis, L., Denis, N.A. St., Vilk, G. and Litchfield, D.W. (2010) Chapter Twenty-Three Protein Kinase CK2 is a Constitutively Active Enzyme that Promotes Cell Survival: Strategies to Identify CK2 Substrates and Manipulate its Activity in Mammalian Cells. Methods Enzymol., 484, 471–493.

9. Halloran, D., Pandit, V. and Nohe, A. (2022) The Role of Protein Kinase CK2 in Development and Disease Progression: A Critical Review. J. Dev. Biol., 10, 31.

10. Stazio, M.D., Zanus, C., Faletra, F., Pesaresi, A., Ziccardi, I., Morgan, A., Girotto, G., Costa, P., Carrozzi, M., d’Adamo, A.P., et al. (2023) Haploinsufficiency as a Foreground Pathomechanism of Poirer-Bienvenu Syndrome and Novel Insights Underlying the Phenotypic Continuum of CSNK2B-Associated Disorders. Genes, 14, 250.

11. Dominguez, I., Cruz-Gamero, J.M., Corasolla, V., Dacher, N., Rangasamy, S., Urbani, A., Narayanan, V. and Rebholz, H. (2021) Okur-Chung neurodevelopmental syndrome-linked CK2α variants have reduced kinase activity. Hum Genet, 140, 1077–1096.

12. Kavaliova, H., Lecis, B., Ballardin, D., Cobret, L., Bienvenu, T., Morisset-Lopez, S. and Rebholz, H. (2025) Pathogenic missense variants of CSNK2B associated with Poirier-Bienvenu neurodevelopmental disorder impact differently on CK2 holoenzyme formation. Biol. Chem., 406, 139–154.

13. Cruz-Gamero, J.M., Ballardin, D., Lecis, B., Zhang, C.-L., Cobret, L., Gast, A., Morisset-Lopez, S., Piskorowski, R., Langui, D., Jose, J., et al. (2025) Missense mutation in the activation segment of the kinase CK2 models Okur-Chung neurodevelopmental disorder and alters the hippocampal glutamatergic synapse. Mol. Psychiatry, 30, 1497–1509.

14. Her, Y., Pascual, D.M., Goldstone-Joubert, Z. and Marcogliese, P.C. (2024) Variant functional assessment in Drosophila by overexpression: what can we learn? Genome, 0, 158–167.

15. Marcogliese, P.C., Deal, S.L., Andrews, J., Harnish, J.M., Bhavana, V.H., Graves, H.K., Jangam, S., Luo, X., Liu, N., Bei, D., et al. (2022) Drosophila functional screening of de novo variants in autism uncovers damaging variants and facilitates discovery of rare neurodevelopmental diseases. Cell Reports, 38, 110517.

16. McGuire, S.E., Mao, Z. and Davis, R.L. (2004) Spatiotemporal Gene Expression Targeting with the TARGET and Gene-Switch Systems in Drosophila. Sci Stke, 2004, pl6.

17. Swarup, S. and Verheyen, E.M. (2012) Wnt/Wingless Signaling in Drosophila. Cold Spring Harb. Perspect. Biol., 4, a007930.

18. Tripathi, B.K. and Irvine, K.D. (2022) The wing imaginal disc. Genetics, 220, iyac020.

19. Owen, C.I., Bowden, R., Parker, M.J., Patterson, J., Patterson, J., Price, S., Sarkar, A., Castle, B., Deshpande, C., Splitt, M., et al. (2018) Extending the phenotype associated with the CSNK2A1-related Okur–Chung syndrome—A clinical study of 11 individuals. Am J Med Genet A, 176, 1108–1114.

20. Wu, R., Tang, W., Liang, L., Li, X., Ouyang, N. and Meng, Z. (2020) Identification of a novel de novo variant of CSNK2A1 gene in a boy with Okur-Chung neurodevelopmental syndrome. Chin J Medical Genetics, 37, 641–644.

21. Chiu, A.T.G., Pei, S.L.C., Mak, C.C.Y., Leung, G.K.C., Yu, M.H.C., Lee, S.L., Vreeburg, M., Pfundt, R., Burgt, I. van der, Kleefstra, T., et al. (2018) Okur-Chung neurodevelopmental syndrome: Eight additional cases with implications on phenotype and genotype expansion. Clin Genet, 93, 880–890.

22. Nakashima, M., Tohyama, J., Nakagawa, E., Watanabe, Y., Siew, C.G., Kwong, C.S., Yamoto, K., Hiraide, T., Fukuda, T., Kaname, T., et al. (2019) Identification of de novo CSNK2A1 and CSNK2B variants in cases of global developmental delay with seizures. J Hum Genet, 64, 313–322.

23. Iossifov, I., O’Roak, B.J., Sanders, S.J., Ronemus, M., Krumm, N., Levy, D., Stessman, H.A., Witherspoon, K.T., Vives, L., Patterson, K.E., et al. (2014) The contribution of de novo coding mutations to autism spectrum disorder. Nature, 515, 216–221.

24. Ernst, M.E., Baugh, E.H., Thomas, A., Bier, L., Lippa, N., Stong, N., Mulhern, M.S., Kushary, S., Akman, C.I., Heinzen, E.L., et al. (2021) CSNK2B: A broad spectrum of neurodevelopmental disability and epilepsy severity. Epilepsia, 62, e103–e109.

25. Li, J., Gao, K., Cai, S., Liu, Y., Wang, Y., Huang, S., Zha, J., Hu, W., Yu, S., Yang, Z., et al. (2019) Germline de novo variants in CSNK2B in Chinese patients with epilepsy. Sci Rep-uk, 9, 17909.

26. Birnbaum, M.J. and Glover, C.V.C. (1991) The phosphotransferase activity of casein kinase II is required for its physiological function in vivo. Biochem. Biophys. Res. Commun., 181, 524–528.

27. Cook, R.K., Christensen, S.J., Deal, J.A., Coburn, R.A., Deal, M.E., Gresens, J.M., Kaufman, T.C. and Cook, K.R. (2012) The generation of chromosomal deletions to provide extensive coverage and subdivision of the Drosophila melanogaster genome. Genome Biol., 13, R21.

28. Nicholson, L., Singh, G.K., Osterwalder, T., Roman, G.W., Davis, R.L. and Keshishian, H. (2008) Spatial and Temporal Control of Gene Expression in Drosophila Using the Inducible GeneSwitch GAL4 System. I. Screen for Larval Nervous System Drivers. Genetics, 178, 215–234.

29. Meggio, F. and Pinna, L.A. (2003) One-thousand-and-one substrates of protein kinase CK2? FASEB J., 17, 349–368.

30. Roffey, S.E. and Litchfield, D.W. (2021) CK2 Regulation: Perspectives in 2021. Biomedicines, 9, 1361.

31. Song, D.H., Dominguez, I., Mizuno, J., Kaut, M., Mohr, S.C. and Seldin, D.C. (2003) CK2 Phosphorylation of the Armadillo Repeat Region of β-Catenin Potentiates Wnt Signaling*. J. Biol. Chem., 278, 24018–24025.

32. Willert, K., Brink, M., Wodarz, A., Varmus, H. and Nusse, R. (1997) Casein kinase 2 associates with and phosphorylates Dishevelled. Embo J, 16, 3089–3096.

33. Wang, S. and Jones, K.A. (2006) CK2 Controls the Recruitment of Wnt Regulators to Target Genes In Vivo. Curr Biol, 16, 2239–2244.

34. Bae, S.M. and Hong, J.Y. (2018) The Wnt Signaling Pathway and Related Therapeutic Drugs in Autism Spectrum Disorder. Clin. Psychopharmacol. Neurosci., 16, 129–135.

35. Kalkman, H.O. (2012) A review of the evidence for the canonical Wnt pathway in autism spectrum disorders. Mol Autism, 3, 10.

36. Walter, N.M., Ohki, C.M.Y., Rickli, M., Smigielski, L., Walitza, S. and Grünblatt, E. (2024) An investigation on the alterations in Wnt signaling in ADHD across developmental stages. Neurosci. Appl., 3, 104070.

37. Pascual, D.M., Rizi, D.J., Kaur, H. and Marcogliese, P.C. (2025) Excess Wnt in neurological disease. Biochem. J., 482, 601–618.

38. Narcisi, R., Arikan, O.H., Lehmann, J., Berge, D. ten and van Osch, G.J.V.M. (2016) Differential Effects of Small Molecule WNT Agonists on the Multilineage Differentiation Capacity of Human Mesenchymal Stem Cells. Tissue Eng. Part A, 22, 1264–1273.

39. Niefind, K., Guerra, B., Ermakowa, I. and Issinger, O. (2001) Crystal structure of human protein kinase CK2: insights into basic properties of the CK2 holoenzyme. EMBO J., 20, 5320–5331.

40. Graham, K.C. and Litchfield, D.W. (2000) The Regulatory β Subunit of Protein Kinase CK2 Mediates Formation of Tetrameric CK2 Complexes*. J. Biol. Chem., 275, 5003–5010.

41. CANTON, D.A., ZHANG, C. and LITCHFIELD, D.W. (2001) Assembly of protein kinase CK2: investigation of complex formation between catalytic and regulatory subunits using a zinc-finger-deficient mutant of CK2β. Biochem. J., 358, 87.

42. Guerra, B. and Issinger, O. (1999) Protein kinase CK2 and its role in cellular proliferation, development and pathology. ELECTROPHORESIS, 20, 391–408.

43. Guerra, B. and Issinger, O.-G. (2020) Role of Protein Kinase CK2 in Aberrant Lipid Metabolism in Cancer. Pharmaceuticals, 13, 292.

44. Guerra, B., Dembic, M., Siddiqui, M.A., Dominguez, I., Ceppi, P. and Andresen, B.S. (2020) Down-Regulation of CK2α Leads toUp-Regulation of the Cyclin-Dependent Kinase Inhibitor p27KIP1 in Conditions Unfavorable for the Growth of Myoblast Cells. Cell. Physiol. Biochem., 54, 1177–1198.

45. Bibby, A.C. and Litchfield, D.W. (2005) The Multiple Personalities of the Regulatory Subunit of Protein Kinase CK2: CK2 Dependent and CK2 Independent Roles Reveal a Secret Identity for CK2β. Int. J. Biol. Sci., 1, 67–79.

46. Gast, A., Schreiber, S. and Jose, J. (2025) Rapid method for evaluation of CK2 enzymatic activity and CK2α/CK2β-interaction in Escherichia coli cell lysates. Biol. Chem., 406, 117–124.

47. Karczewski, K.J., Francioli, L.C., Tiao, G., Cummings, B.B., Alföldi, J., Wang, Q., Collins, R.L., Laricchia, K.M., Ganna, A., Birnbaum, D.P., et al. (2020) The mutational constraint spectrum quantified from variation in 141,456 humans. Nature, 581, 434–443.

48. Ballardin, D., Cruz-Gamero, J.M., Bienvenu, T. and Rebholz, H. (2022) Comparing Two Neurodevelopmental Disorders Linked to CK2: Okur-Chung Neurodevelopmental Syndrome and Poirier-Bienvenu Neurodevelopmental Syndrome—Two Sides of the Same Coin? Front. Mol. Biosci., 9, 850559.

49. Lozeman, F.J., Litchfield, D.W., Piening, C., Takio, K., Walsh, K.A. and Krebs, E.G. (1990) Isolation and characterization of human cDNA clones encoding the .alpha. and the .alpha.’ subunits of casein kinase II. Biochemistry, 29, 8436–8447.

50. Unni, P., Friend, J., Weinberg, J., Okur, V., Hochscherf, J. and Dominguez, I. (2022) Predictive functional, statistical and structural analysis of CSNK2A1 and CSNK2B variants linked to neurodevelopmental diseases. Front. Mol. Biosci., 9, 851547.

51. Gupta-Rossi, N., Ortica, S., Meas-Yedid, V., Heuss, S., Moretti, J., Olivo-Marin, J.-C. and Israël, A. (2011) The Adaptor-associated Kinase 1, AAK1, Is a Positive Regulator of the Notch Pathway*. J. Biol. Chem., 286, 18720–18730.

52. Baonza, A. and Garcia-Bellido, A. (2000) Notch signaling directly controls cell proliferation in the Drosophila wing disc. Proc. Natl. Acad. Sci., 97, 2609–2614.

53. Blanc, A., Bonnet, C., Wandzel, M., Roth, V., Duffourd, Y., Safraou, H., Leheup, B., Muller, F., Colne, J.D., Feillet, F., et al. (2024) Patient with a heterozygous pathogenic variant in CSNK2A1 gene: A new case to update the Okur–Chung neurodevelopmental syndrome. Am. J. Méd. Genet. Part A, 194, e63642.

54. Ranganath, P., Ranganath, P., Vineeth, V.S., Dalal, A. and Patil, S.J. (2021) Report of an Asian–Indian patient with Okur–Chung Syndrome and comparison of the clinical phenotype in different ethnic groups. Clin. Dysmorphol., 30, 209–212.

55. Hassett, K., Potu, S.S., Sankaramoorthy, A., Kaneshamoorthy, S., Leka, K., Huentelman, M.J., Narayanan, V. and Rangasamy, S. (2024) Characterizing CSNK2A1 Mutant-Induced Morphological Phenotypes in Zebrafish (Danio rerio): Insights into Okur-Chung Neurodevelopmental Syndrome (OCNDS). bioRxiv, 2024.01.09.574075.

56. Lou, D.Y., Dominguez, I., Toselli, P., Landesman-Bollag, E., O’Brien, C. and Seldin, D.C. (2008) The Alpha Catalytic Subunit of Protein Kinase CK2 Is Required for Mouse Embryonic Development. Mol Cell Biol, 28, 131–139.

57. Buchou, T., Vernet, M., Blond, O., Jensen, H.H., Pointu, H., Olsen, B.B., Cochet, C., Issinger, O.-G. and Boldyreff, B. (2003) Disruption of the regulatory beta subunit of protein kinase CK2 in mice leads to a cell-autonomous defect and early embryonic lethality. Mol Cell Biol, 23, 908–15.

58. Padmanabha, R., Chen-Wu, J.L.-P., Hanna, D.E. and Glover, C.V.C. (1990) Isolation, Sequencing, and Disruption of the Yeast CKA2 Gene: Casein Kinase II Is Essential for Viability in Saccharomyces cerevisiae. Mol. Cell. Biol., 10, 4089–4099.

59. Zhang, S., Gong, S., Yang, Y., Gong, W., Zhou, Q., Yang, K., Rui, M. and Wang, S. (2026) Casein kinase 2 maintains the self-renewal of neural stem cells via prospero phosphorylation in Drosophila. Cell Biosci., 16, 23.

60. Bagatelas, E.D., Khan, M.M. and Rushing, G.V. (2025) OCNDS core features are conserved across variants, with loop-region mutations driving greater symptom burden. Front. Hum. Neurosci., 19, 1589897.

61. Mulligan, K.A. and Cheyette, B.N.R. (2017) Neurodevelopmental Perspectives on Wnt Signaling in Psychiatry. Mol. Neuropsychiatry, 2, 219–246.

62. Patapoutian, A. and Reichardt, L.F. (2000) Roles of Wnt proteins in neural development and maintenance. Curr. Opin. Neurobiol., 10, 392–399.

63. Chai, G., Szenker-Ravi, E., Chung, C., Li, Z., Wang, L., Khatoo, M., Marshall, T., Jiang, N., Yang, X., McEvoy-Venneri, J., et al. (2021) A Human Pleiotropic Multiorgan Condition Caused by Deficient Wnt Secretion. N. Engl. J. Med., 385, 1292–1301.

64. Martin, P.-M., Yang, X., Robin, N., Lam, E., Rabinowitz, J.S., Erdman, C.A., Quinn, J., Weiss, L.A., Hamilton, S.P., Kwok, P.-Y., et al. (2013) A rare WNT1 missense variant overrepresented in ASD leads to increased Wnt signal pathway activation. Transl. Psychiatry, 3, e301–e301.

65. Kuechler, A., Willemsen, M.H., Albrecht, B., Bacino, C.A., Bartholomew, D.W., van Bokhoven, H., van den Boogaard, M.J.H., Bramswig, N., Büttner, C., Cremer, K., et al. (2015) De novo mutations in beta-catenin (CTNNB1) appear to be a frequent cause of intellectual disability: expanding the mutational and clinical spectrum. Hum. Genet., 134, 97–109.

66. Asif, M., Kaygusuz, E., Shinawi, M., Nickelsen, A., Hsieh, T.-C., Wagle, P., Budde, B.S., Hochscherf, J., Abdullah, U., Höning, S., et al. (2022) De novo variants of CSNK2B cause a new intellectual disability-craniodigital syndrome by disrupting the canonical Wnt signaling pathway. Hum. Genet. Genom. Adv., 3, 100111.

67. Landrum, M.J., Chitipiralla, S., Brown, G.R., Chen, C., Gu, B., Hart, J., Hoffman, D., Jang, W., Kaur, K., Liu, C., et al. (2019) ClinVar: improvements to accessing data. Nucleic Acids Res, 48, D835–D844.

68. Calleja, M., Herranz, H., Estella, C., Casal, J., Lawrence, P., Simpson, P. and Morata, G. (2000) Generation of medial and lateral dorsal body domains by the pannier gene of Drosophila. *Development (Cambridge*, England), 127, 3971–3980.

69. Ma, M., Zhang, X., Zheng, Y., Lu, S., Pan, X., Mao, X., Pan, H., Chung, H., Wang, H., Guo, H., et al. (2022) The fly homolog of SUPT16H, a gene associated with neurodevelopmental disorders, is required in a cell-autonomous fashion for cell survival. Hum. Mol. Genet., 32, 984–997.

70. Dutta, D., Kanca, O., Shridharan, R.V., Marcogliese, P.C., Steger, B., Morimoto, M., Frost, F.G., Macnamara, E., Network, U.D., Wangler, M.F., et al. (2024) Loss of the endoplasmic reticulum protein Tmem208 affects cell polarity, development, and viability. Proc. Natl. Acad. Sci., 121, e2322582121.

71. Bischof, J., Björklund, M., Furger, E., Schertel, C., Taipale, J. and Basler, K. (2013) A versatile platform for creating a comprehensive UAS-ORFeome library in Drosophila. Development, 140, 2434–2442.

72. Venken, K.J.T., He, Y., Hoskins, R.A. and Bellen, H.J. (2006) P[acman]: a BAC transgenic platform for targeted insertion of large DNA fragments in D. melanogaster. *Science (New York*, N.Y.), 314, 1747–1751.

73. Marcogliese, P.C., Dutta, D., Ray, S.S., Dang, N.D.P., Zuo, Z., Wang, Y., Lu, D., Fazal, F., Ravenscroft, T.A., Chung, H., et al. (2022) Loss of IRF2BPL impairs neuronal maintenance through excess Wnt signaling. Sci Adv, 8, eabl5613.

74. Accogli, A., Lu, S., Musante, I., Scudieri, P., Rosenfeld, J.A., Severino, M., Baldassari, S., Iacomino, M., Riva, A., Balagura, G., et al. (2022) Loss of Neuron Navigator 2 Impairs Brain and Cerebellar Development. Cerebellum, 1–17.

75. Goodman, L.D., Cope, H., Nil, Z., Ravenscroft, T.A., Charng, W.-L., Lu, S., Tien, A.-C., Pfundt, R., Koolen, D.A., Haaxma, C.A., et al. (2021) TNPO2 variants associate with human developmental delays, neurologic deficits, and dysmorphic features and alter TNPO2 activity in Drosophila. Am J Hum Genetics.

76. Ioannidis, N.M., Rothstein, J.H., Pejaver, V., Middha, S., McDonnell, S.K., Baheti, S., Musolf, A., Li, Q., Holzinger, E., Karyadi, D., et al. (2016) REVEL: An Ensemble Method for Predicting the Pathogenicity of Rare Missense Variants. Am. J. Hum. Genet., 99, 877–885.

77. Dong, C., Wei, P., Jian, X., Gibbs, R., Boerwinkle, E., Wang, K. and Liu, X. (2015) Comparison and integration of deleteriousness prediction methods for nonsynonymous SNVs in whole exome sequencing studies. Hum. Mol. Genet., 24, 2125–2137.

