## Supplementary figures and images for "CK2 variant function and disease modelling in *Drosophila* reveal allelic heterogeneity and Wnt/β-catenin-mediated phenotypes"

### Supplementary Material, Fig. S1

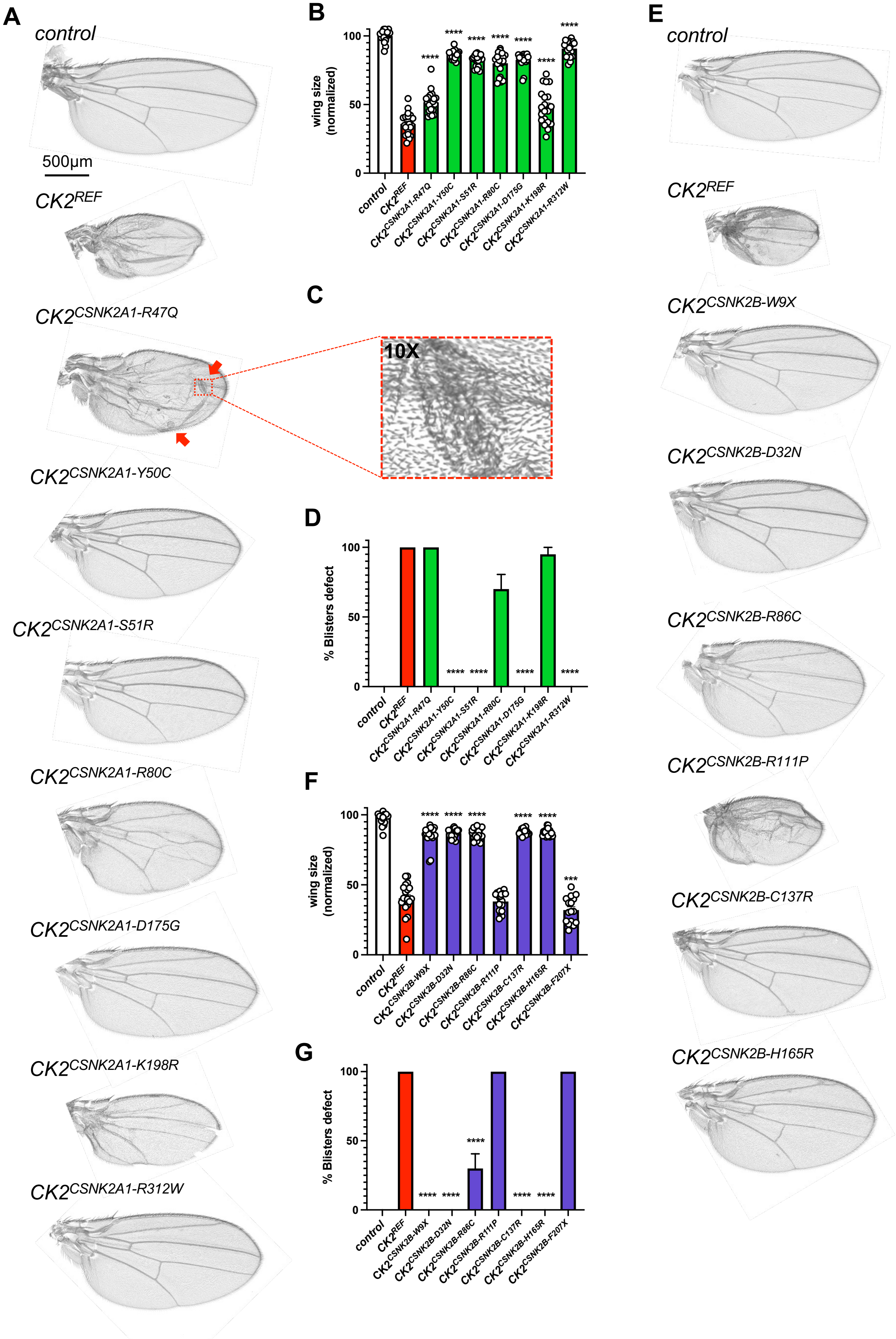

### Supplementary Material, Fig. S2

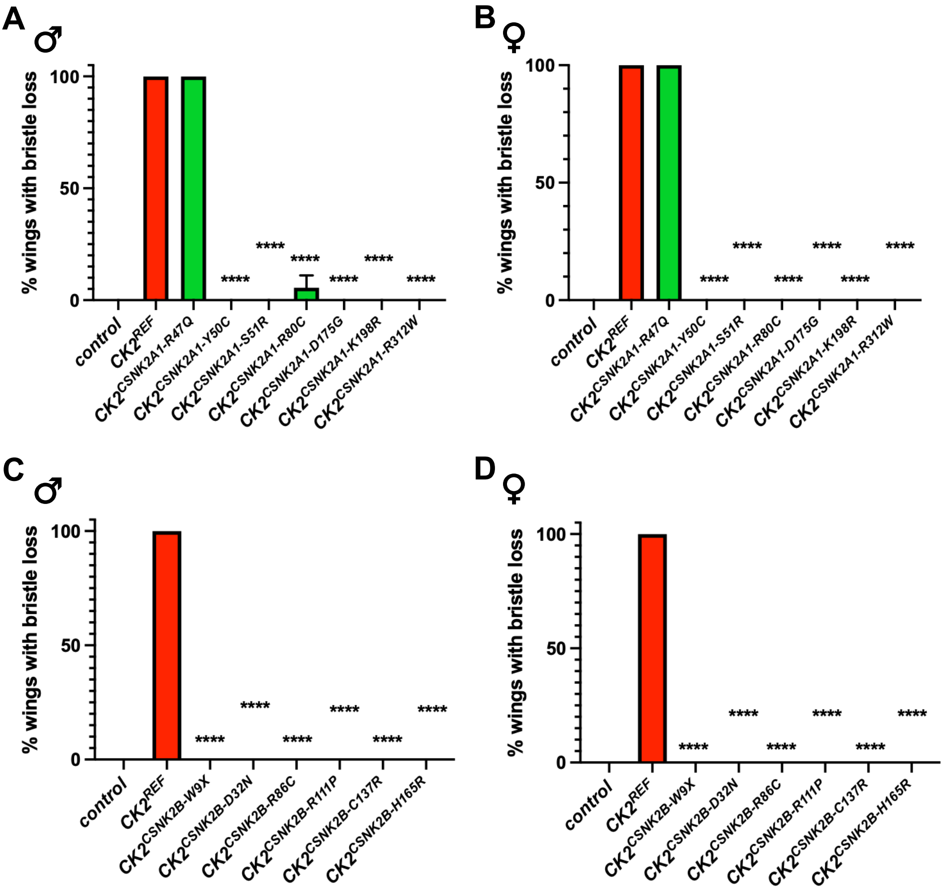

### Supplementary Material, Fig. S3

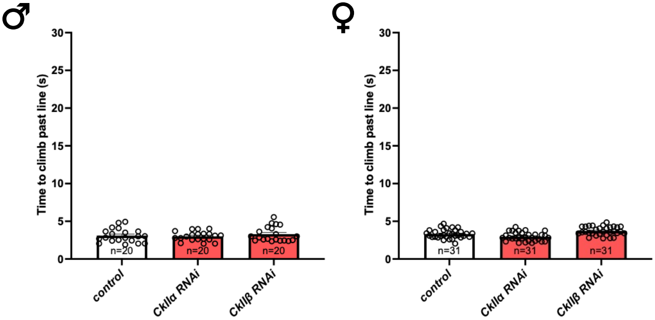
